# Fluorinated RNA origami enables serum-stable nanodevices for sensing and targeting

**DOI:** 10.64898/2026.03.04.709525

**Authors:** Emil Laust Kristoffersen, Nikolaj Holck Zwergius, Nestor Sampedro Vallina, Amanda Dyrholm Stange, Lasse Messell Desdorf, Sofie Thomsen, Victoria Birkedal, Nicolas Glück, Laia Civit, Cody Geary, Jørgen Kjems, Julian Valero, Ebbe Sloth Andersen

## Abstract

Chemically modified RNAs with increased stability and reduced immunogenicity have transformed RNA therapeutics. Rational RNA design methods, including RNA origami, seek to further extend RNA medicine and biotechnology by encoding advanced functions such as signalling, targeting, and controlled release within the RNA polymer. However, current design methods lack the ability to integrate chemical modification or predict how it shapes the structure of large RNA assemblies inhibiting its use in RNA therapeutics. Here we demonstrate that 2’-fluoro pyrimidine RNA (FY-RNA) origami structures can be co-transcriptionally folded to generate serum-stable nanodevices. Cryogenic electron microscopy reveals that FY-RNA can alter folding pathways and perturb tertiary motifs, while molecular dynamics simulations show how 2’-fluoro modification affects hydrogen bonding, sugar pucker, and helix-helix interactions. Despite these structural perturbations, fluorogenic aptamers embedded within RNA origami retain partial activity and enable logic-based molecular sensing in human serum. Finally, we use an FY-RNA scaffold to determine the structure of an FY-RNA anti-Spike aptamer bound to the Spike protein at 3.4 Å resolution, uncovering fluorine-specific structural motifs and protein interactions. Together, our results establish design principles for nuclease-resistant RNA architectures and position FY-RNA as a versatile polymer for constructing medical nanodevices and environmental sensors. More broadly, this work provides a framework for systematically exploring the folding landscape of chemically modified RNAs, expanding the chemical and functional diversity accessible to nucleic acid nanotechnology and RNA medicine.

## Introduction

The ability of nucleic acids to encode information while adopting defined three-dimensional folds underpins both biology and biotechnology^1,2^. Beyond canonical DNA and RNA, nature employs more than 170 chemical modifications that expand the chemical and functional repertoire of nucleic acids^3^. Synthetic analogues, collectively termed xenonucleic acids (XNAs)^4-8^, further extend this diversity and can be chemically synthesized or enzymatically synthesized using engineered polymerases^5,9^. The ability to synthesize XNAs have been widely used to improve RNA medicine - from the stabilization of siRNA drugs^10^ to decreasing the immunogenicity of mRNA vaccines^11,12^. Furthermore, it has been shown that XNAs have special folding properties^13^ and makes both functional^6,14-21^ and catalytic folds^12,18,19^. Consequently XNAs have emerged as powerful platforms in biotechnology and therapeutics, where chemical diversification enhances stability, modulates immunogenicity, and preserves or expands biological function^10-12,22^.

Rational design of nucleic acids has been another important development contributing new functionalities for medicine and biotechnology. Compared to protein design^23^, nucleic acids have traditionally been easier to design because of the simple base pairing rules and well defined double helix^24^. The DNA origami method^25^ provided a general approach to design programmable architectures that self-assemble by heat-annealing. Because this method relies on chemical synthesis of DNA strands it is very easy to add chemical functionality which has been used to improve properties for drug delivery^26^. The RNA origami method^27^ was developed for enzymatic production of nanostructures and takes advantage of the co-transcriptional folding and unique structural motifs available for RNA. However, unlike DNA origami, chemical modification has played only a limited role in RNA origami, as incorporation must occur during enzymatic synthesis or through post-transcriptional modification^28^. Initial efforts have demonstrated the usefulness of various XNAs for designing nanostructures^29,30^ but the enzymatic production and co-transcriptional folding of XNA origami has not been widely explored.

2’-fluoro pyrimidine RNA (FY-RNA) is a well-characterized XNA (Fig. 1a), which forms a regular A-form double helix^31^, but is distinguished by improved thermal stability^16,32^, extreme serum stability^16^, altered immunogenicity^33^, and high biocompatibility^10^. Further, the fluorine (F) atom at the 2’-position of the ribose sugar gives the polymer new properties based on increased base pairing and stacking stability^31,34^, altered hydrogen bonds (H-bonds) where the F atom acts as an acceptor with modest strength^35^ (Fig. 1b), sugar conformation favouring C3’-endo^36^ (Fig. 1c), and hydrophobicity of the shallow groove where F makes it more hydrophobic^36^. FY-RNA strands can be synthesized efficiently via *in vitro* transcription using a mutant RNA polymerase^37^, which has allowed *in vitro* selection of several FY-RNA aptamers^14,15^,38,39. FY-RNA transcription in principle allows co-transcriptional folding of single-stranded FY-RNA origami (Fig. 1d) and has been used to develop an FY-RNA origami-based anticoagulant device^40,41^ that functions *in vivo* to reduce blood coagulation in mice^42^. However, FY-RNA origamis have not been structurally characterized and the effects of 2’-F-modification on structural motifs and folding pathways remain to be studied.

**Figure 1.**
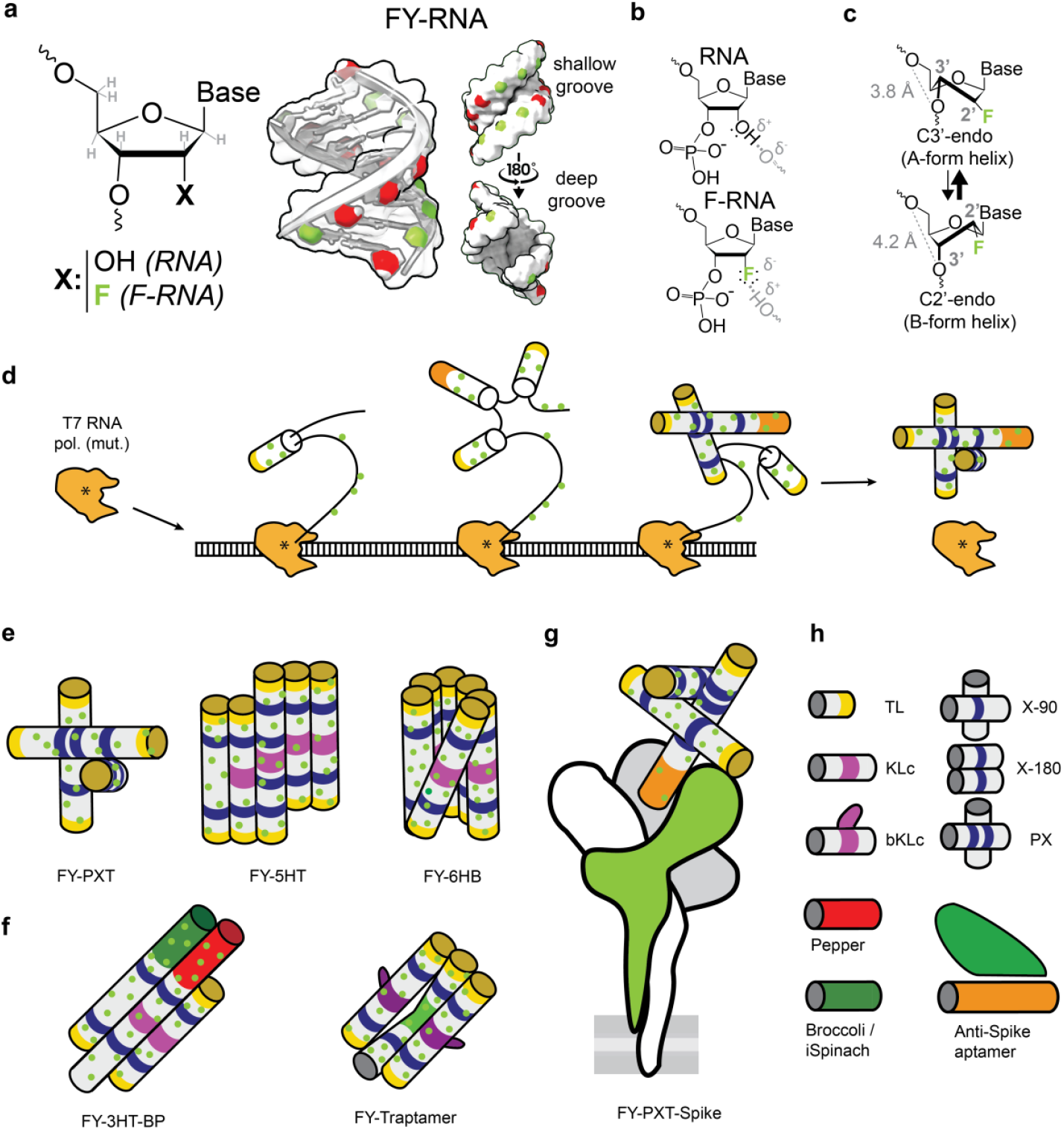
Co-transcriptional folding of FY-RNA origami nanostructures. **a**, Chemical structure of ribose with either 2’-F modification or 2’-OH and the FY-RNA A-form helix which presents the 2’-F atoms in the shallow groove leaving the deep groove unchanged. F is coloured light green according to the CPK colouring convention. **b**, The highly electronegative F atom in F-RNA with three lone pairs subtracts electrons from its local environment in the molecule (illustrated with δ^+^ and δ^−^) and is a weak H-bond acceptor. In contrast, the hydroxyl (OH) in canonical RNA is an H-bond donor. **c**, The F atom stabilizes the C3’-endo conformation compared to RNA. **d**, Cartoon illustration of co-transcriptional production of single-stranded FY-RNA origami using the mutant (Y639F) T7 RNA polymerase (mutation indicated by *). **e-g**, Cylinder models of FY-RNA scaffolds: PXT, 5HT, and 6HB, and FY-RNA devices: 3HT-BP, Traptamer, and PXT-Spike ‘pointer’ bound to the Spike protein of SARS-CoV-2. **h**, FY-RNA modules: Tetra loop (TL, yellow), 90 degree crossover (X-90, blue), paranemic crossover (PX, blue), 180 degree crossover (X-180, blue), kissing loop (KL, purple), branched KL (bKL, purple), fluorogenic aptamers Pepper and Broccoli/iSpinach (red and green, respectively); anti-Spike binding aptamer (orange) binding the RBD (light green). All models are coloured by motifs. Green spots depict the 2’-F.

Here we present a detailed investigation of the structure, folding and function of FY-RNA origamis using cryo-EM characterization and molecular dynamics (MD) simulation. Based on these structural studies we demonstrate large, highly ordered, and rationally designed FY-RNA nanostructures with advanced functions that are stable and functional in human serum. Together this work establishes FY-RNA origami as a generalizable technique for engineering advanced serum-stable nanodevices and pinpoints precise structural effects and design principles of 2’-F-modification within a macromolecular context.

## Results

### Platform for characterizing FY-RNA structures

In this study we investigate a panel of FY-RNA origami scaffolds and devices (Fig. 1e-g) with the aim of understanding the effects of 2’-FY modification on structural and functional motifs (Fig. 1h). To study the effect on structural motifs, we selected a paranemic crossover triangle (PXT)^43^, a 5-helix tile (5HT)^44^, and a 6-helix bundle (6HB)^44^, which contains A-form helices (H), tetraloops (TL), crossovers (X), and kissing loops (KL). To study the effect on functional motifs, we selected a 3-helix tile that contains the fluorogenic aptamers Broccoli and Pepper (3HT-BP)^45^ and a recent RNA origami robot, named the Traptamer^46^, that contains the fluorogenic aptamer iSpinach and two branched kissing loops (bKL). Since these RNA origami structures have been previously characterized, we can directly compare them to the FY-RNA version made in this study. Finally, we designed a novel “pointer” device to facilitate the structure determination of an FY-RNA aptamer binding to the receptor binding domain (RBD) of the SARS-CoV-2 Spike protein^14^ (PXT-Spike, Fig. 1g). Blueprints and sequences for the RNA origamis used are found in Fig. S1-7 and Table S1 (see supplementary material).

FY-RNA transcriptions of the RNA origami templates yielded full-length and homogeneous products that were both resistant to nucleases and stable in human serum (Fig. E1, see supplementary material). The FY-RNA nanostructures were studied by cryo-EM single particle analysis to obtain electrostatic potential maps at resolutions of 2.9-10 Å (Fig. S8-14 and Table S2-5). Since current modelling software does not support unnatural chemically modifications of RNA structures, we developed a software named Nucleic Acid Modelling including Xenonucleotides (NAMiX) to solve this issue (Materials and Methods, Data availability, Fig. E2). NAMiX facilitated the construction of atomic models of the FY-RNA origamis (Fig. E3), which were investigated in further detail by molecular dynamics (MD) simulation using a recently published force field for the 2’-F modification^47^ (Data availability, Fig. S15-21). The combined use of cryo-EM, NAMiX, MD, and functional assays provides a powerful platform for studying how natural and synthetic RNA modifications influence structure and function.

### 2’-FY modification affects RNA origami structure and folding

The cryo-EM structure of the FY-RNA PXT origami was determined to a resolution of 4.4 Å (Fig. 2a, Fig. S8) and the resulting FY-RNA model (PDB: 9R7Q) was found to be very similar to the previously reported RNA version (PDB: 8BTZ^43^) as indicated by a root-mean square deviation (RMSD) of 3.3 Å (Fig. E4a). The PXT contains six tetraloops (TL) with the loop sequence UUCG, that adopts a stable tertiary structure as RNA^48^ including two H-bonds and two nucleotides in C2’-endo conformation which will be affected as FY-RNA. The TLs have poor resolution, since they are placed at helix ends, which is the most flexible part of the structure. The best resolved TL was TL3 with a local resolution of ∼6-7 Å (Fig. 2b) which does not allow observation of differences between RNA and FY-RNA TLs. This prompted us to investigate the TLs by MD simulation (see next section). The PXT contains crossovers that are resolved to ∼4 Å: Two single crossovers at 70° and 85° angles between helices (Fig. 2c) and a paranemic crossover (PX) composed of two 85° crossovers at a spacing of only six base pairs across the deep groove^43^ (Fig. 2d). The crossovers have four central nucleotides N1-N4, where N1 and N3 are often found with C2’-endo sugar pucker^44^. The FY-RNA PXT has five examples where N1 or N3 are FY, which could negatively affect the crossover because of the reported preference of 2’-F modified RNA for C3’-endo sugar pucker^36^. However, no major differences are observed between the RNA and FY-RNA maps. This suggests that N1 and N3 of the FY-RNA crossovers are adopting the C2’-endo conformation, which is further supported by MD simulations (see next section).

**Figure 2.**
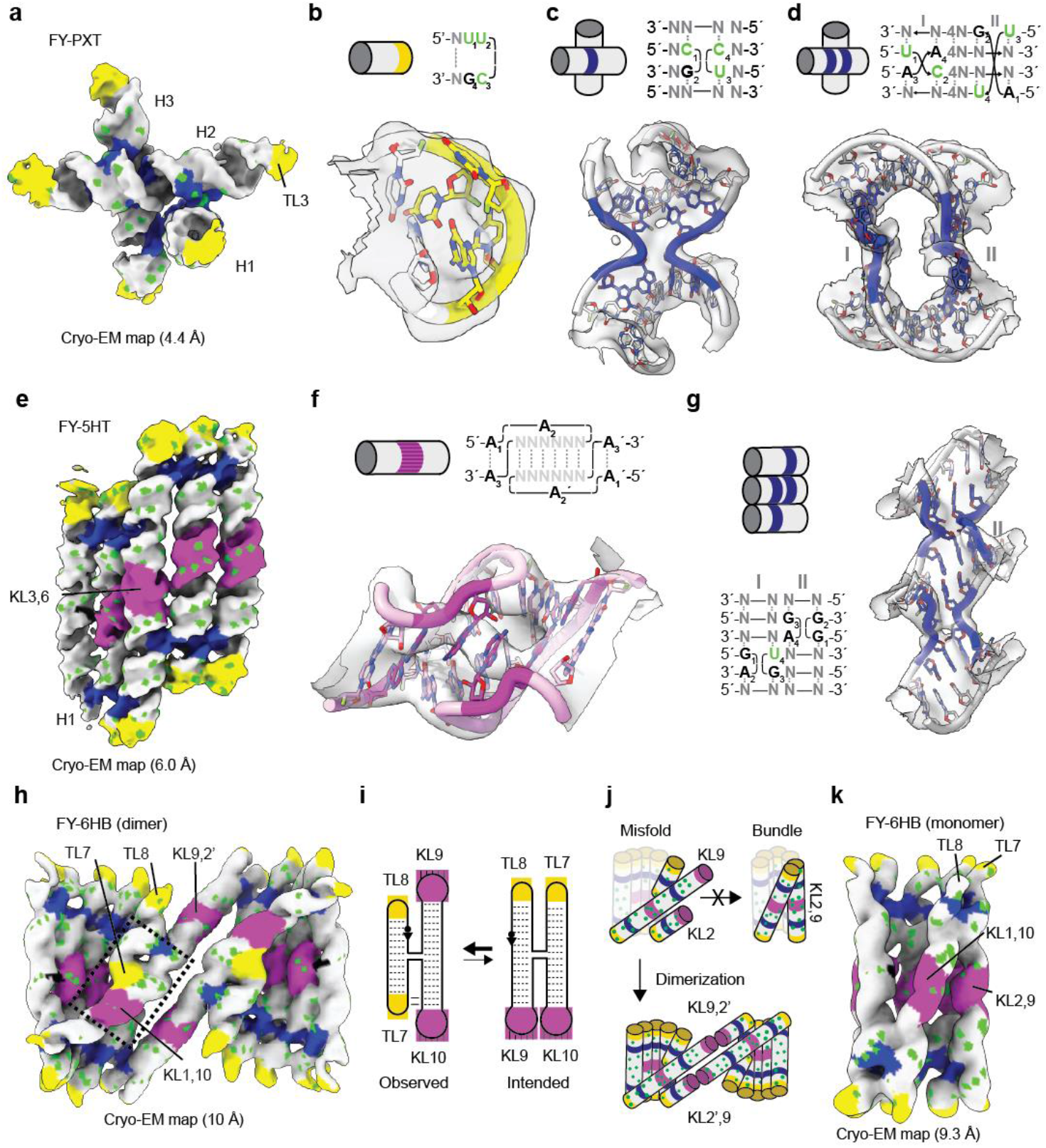
Cryo-EM of FY-RNA origami scaffolds and structural motifs. **a**, Cryo-EM map of FY-RNA PXT origami at 4.4 Å resolution. **b-d**, PXT TL, X-90, and PX motifs shown as secondary structure diagram and atomic model with overlay of cryo-EM map. **b**, Cryo-EM map of FY-RNA 5HT origami at 6.0 Å resolution. **f**,**g**, 5HT KL and DT motifs shown as secondary structure diagrams and atomic models in cryo-EM map. **h**, Cryo-EM map of FY-RNA 6HB dimer at 10 Å resolution. **i**, Illustration of the observed (left) and intended (right) crossover arrangements. **j**, Misfolded structure leads to dimerization and inhibits folding of the intended bundle structure. **k**, New design that leads to formation of the FY-RNA 6HB monomer. Motifs are coloured like in Fig. 1. Green spots depict the 2’-F.

The cryo-EM structure of the FY-RNA 5HT origami was determined to a resolution of 6.0 Å (Fig. 2e, Fig. S9) and the resulting model (PDB: 9R7W) was shown to be similar to the RNA version (PDB: 7QDU^44^) with an RMSD of 5.8 Å (Fig. E4b). The biggest difference is seen in H1 that has an RMSD of up to 20 Å (Fig. E4b). This region has a poor local resolution in both RNA and FY-RNA, indicating flexibility or misfolding (Fig. S9c). The 5HT has four KLs with the loop sequence A1A2(6N)A3, where 6N refers to the six base pairs that form between complementary KLs to make a KL complex^49^. The best resolved KL complex is KL3,6 with a local resolution of ∼6-7 Å (Fig. 2f). The cryo-EM map confirms the tertiary structure of the FY-RNA KL complex including the stacking of A2 and A2’ bases across the deep groove as previously reported for RNA KLs^44^. The 5HT contains eight crossovers at ∼180° angles (X-180) that are arranged in four double crossovers (DX) and six dovetails (DT)^50^. A well-resolved DT is found between H3, H4, and H5 (Fig. 2g). The FY-RNA 5HT only has one crossover where N3 is FY, which will likely adopt the C2’-endo sugar pucker as supported by MD simulations (see next section). In conclusion, we find that KLs and 180° crossovers can be directly converted and used as structural modules in FY-RNA origami.

The RNA 6HB origami has previously been shown to possess an interesting folding and maturation path with a young and a mature conformation^44,51^. The cryo-EM structure of the FY-RNA 6HB was determined to a resolution of 10.0 Å and revealed an unexpected homodimeric structure (Fig. 2h). The cryo-EM map was improved to 8.9 Å by C2-symmetry expansion (Fig. S10) and allowed the overall construction of a model of the FY-RNA 6HB dimer (PDB: 9SD9) and a revised secondary structure map (Fig. E5). The model of the dimer shows that each of the monomers resemble the less compact young conformation (PDB: 7PTK^44^) but has an alternative stacking conformation of the crossover between H5 and H6, which results in KL9 facing outwards (Fig. 2i, left) instead of inwards as intended (Fig. 2i, right). This alternative conformation is further stabilized by interactions between TL7 and the shallow groove of the KL10 helix (Fig. 2h, box). The alternative folding pathway arises because the intramolecular KL2,9 fails to form in the FY-RNA context. This disruption propagates downstream, allowing KL9 and KL10 to engage in a stable stacking interaction. Subsequent formation of KL1,10 exposes KL9, which then promotes intermolecular dimerization through pairing with KL2’ (Fig. 2j). Interestingly, similar dimer conformations have been observed when base-modifications are introduced into the 6HB^52^. To improve the folding of the 6HB, we redesigned the sequence and twist-corrected the KL-containing helices^44,50^, which resulted in a monomeric bundle resolved to 9.3 Å by cryo-EM (Fig. 2k, Fig. S10). In summary, we find that 2’-FY modification has minor effects on the folding of RNA origami structures, but that the minor effects can cause major misfolding in more complex designs.

### 2’-FY destabilize RNA structural motifs and helix-helix interactions

All-atom molecular dynamics (MD) simulation was used to investigate how 2’-F modification alters local molecular interactions and conformational flexibility, showing its specific impact on individual structural motifs. To compare FY-RNA and RNA simulations we calculated root-mean square fluctuation (RMSF) of the last 200 ns of the simulations (Fig. E6) and evaluated the difference as ΔRMSF = RMSF_RNA_-RMSF_FY-RNA_ (Fig. S15-18). The core triangulated crossovers in the PXT have similar flexibility in RNA and FY-RNA (Fig. 3a, white). The TL4 and TL6 helices are more flexible as RNA (Fig. 3a, red), which can be explained by the higher rigidity of FY-RNA double helices^31,34^. TL1 and TL3 helices are more flexible as FY-RNA (Fig. 3a, blue), which was caused by stabilizing helix-helix and TL-TL interactions in RNA (Fig. S15 and Video 1). In the FY-RNA PXT the G4 nucleotide was found to be more flexible in both TL1, TL2, TL3 and TL5 (Fig. 3a, blue). The simulation of the 5HT showed that the FY-RNA version is more flexible at helix-helix interfaces, crossovers, and at most TLs (Fig. 3b, Fig. S16-17, and Supplementary Video 2). In conclusion, the UUCG TLs are destabilized and could be replaced by the purine-rich GNRA^64^ TLs in future RNA origami designs.

**Figure 3.**
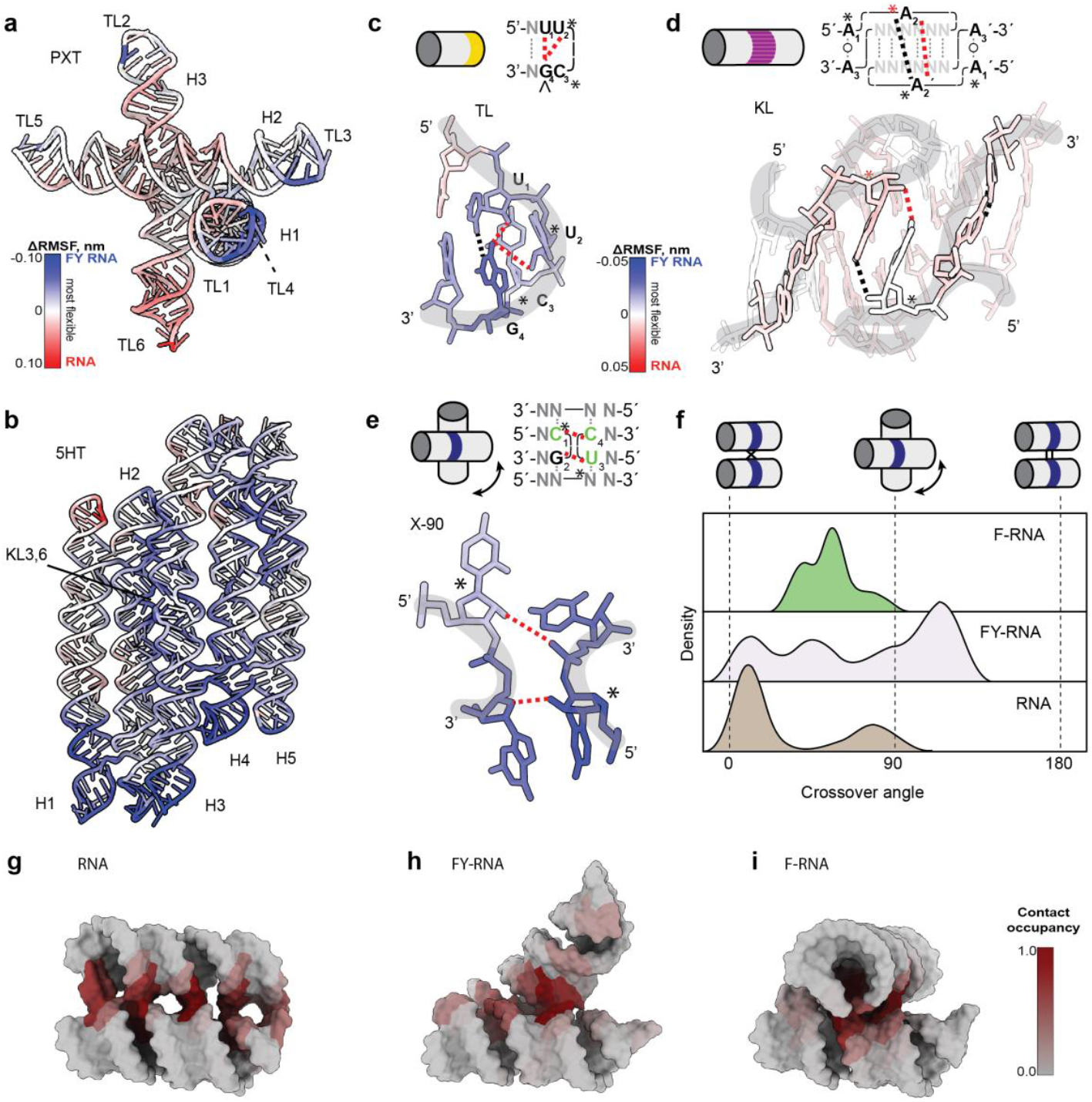
Molecular dynamics simulation of FY-RNA structures and motifs. **a-b**, MD simulation of PXT and 5HT as RNA and FY-RNA coloured by difference in root-mean-square fluctuation (ΔRMSF) as indicated in the scalebar next to the PXT. **c-e**, Secondary structure diagrams and MD simulations of TL, KL and X motifs coloured by ΔRMSF as indicated in the scalebar next to the TL. Dashed black lines mark identified H-bonds whereas the dashed red lines indicate H-bonds that are not observed in the fluorinated RNA. Asterisk marks residues that in the simulations often are observed in C2’-endo conformation. **f**, Angle density plot of MD simulations of X motif as F-RNA, FY-RNA and RNA. **g-i**, RNA, FY-RNA and F-RNA models from MD simulations with contact occupancy plotted on the structures. For full MD simulation data see ‘Data availability’.

Next, we simulated individual motifs as RNA, FY-RNA and fully 2’-F-modified RNA (F-RNA) to investigate the effects of 2’-F modification outside the context of the origami scaffolds. The A-form helix is generally unaffected by the 2’-F modifications as previously reported^31^ (Fig. S18). The FY-RNA TLs show an increased flexibility at the G4 position (Fig. 3c, Fig. S19a), which is caused by the loss of two H-bonds between G4 O6 and the 2’-OHs of U1 and U2^48^. When simulating the KL complex as RNA and FY-RNA, we found that they had very similar RMSF (Fig. 3d, Fig. S19b), indicating that the KL complex is not destabilized as FY-RNA. To investigate the role of A1-3 in the KL, we simulated a FY-RNA KL where A1-3 and A1-3’ were also 2’-F modified. This simulation shows that A2 and A2’ display greater flexibility as 2’-F and that A2 is more prone to shift sugar pucker from C2’-endo to C3’-endo (Fig. S19b)^55^. We then simulated a single crossover from the PXT (between H1 and H3) as both RNA, FY-RNA and F-RNA. We find that the RNA crossover nucleotides N1 and N3 are predominantly in the C2’-endo conformation and form cross-junction H-bonds, consistent with previous findings^44,53^. For FY-RNA and F-RNA the cross-junction H-bonds are partly or fully lost (Fig. 3e) but we surprisingly find that the N1 and N3 positions have a stronger preference for the C2’-endo than RNA (Fig. S20). Thus, the simulations show that the reported preference of F-RNA for C3’-endo conformation^36^ depends on the structural context.

By measuring the crossover angle over the time-course of the simulation, we observe that the RNA crossover primarily arranges to form a ∼0° angle (a parallel crossover), the FY-RNA crossover arranges to many different angles, and the F-RNA crossover favours a ∼45° angle (Fig. 3f, Fig. S20, and Video 3). Analysis of the RNA simulation shows that the 0° crossover is stabilized by the formation of multiple helix-helix interactions formed primarily by H-bonds between 2’-OHs and the phosphodiester backbone (Fig. 3g). The FY-RNA simulation shows reduced 2’-OH interactions along the helices (Fig. 3h), whereas the F-RNA simulation shows that the 45° angle is stabilized by hydrophobic interactions between two shallow grooves next to the crossover (Fig. 3i). The latter observation can be explained by the hydrophobicity of the shallow groove of 2’-F-modified helices^36^. Indeed, these helix-helix interactions explain both simulated and experimental effects in the full origami structures: (1) The PXT RNA simulation showed interactions between the helices of TL1 and TL3 caused by 2’-OH helix-helix interactions. (2) The 5HT FY-RNA simulation showed increased flexibility between parallel helices caused by the lack of 2’-OH helix-helix interactions. (3) The FY-RNA 6HB misfolding is likely affected by the lack of 2’-OH helix-helix interactions that would help parallelize helices and favour the intended mature conformation. Our data highlights the role of helix-helix interactions in promoting the folding and stabilization of RNA origami with parallel helices. In conclusion, the combined MD simulations show that FY-RNA destabilize UUCG TLs, does not affect KLs, destabilize crossovers, and destabilize parallel helix-helix interactions. These insights can be used for rational design of FY-RNA origami structures, where pyrimidines can be avoided at critical positions.

### FY-RNA fluorogenic aptamers with improved serum stability

To study the impact of 2’-F modifications in structurally complex functional motifs, we used the 3HT-BP origami device^45^, which contains the fluorogenic aptamers Broccoli and Pepper that bind the fluorophores HBC620 and DFHBI-1T, respectively^54-58^. The 3HT-BP device places the aptamers in proximity allowing Förster resonance energy transfer (FRET) to occur (Fig. 4a). The FY-RNA 3HT-BP was shown to maintain a FRET efficiency of E ∼ 0.6 (Fig. 4b, Fig. E7a) but also to have reduced fluorescence of both aptamers (Fig. 4d,e) indicating that while the aptamers are negatively affected by 2’-F modification, the conformation of the device is maintained. To investigate these effects further, we used cryo-EM to determine the FY-RNA 3HT-BP structure in the presence of both fluorophores to a resolution of 6.4 Å (locally to 5.5 Å) (Fig. 4c, Fig. S11). The FY-RNA map (EMD-55626) resembles that of RNA (EMD-14740^45^) but differs in the Broccoli region of the map. 3D classification of the FY-RNA cryo-EM particles used in the final map shows that Pepper is present in all classes, whereas Broccoli is missing in some (Fig. S11g), indicating that Broccoli is only folded correctly in a subpopulation of the particles.

**Figure 4.**
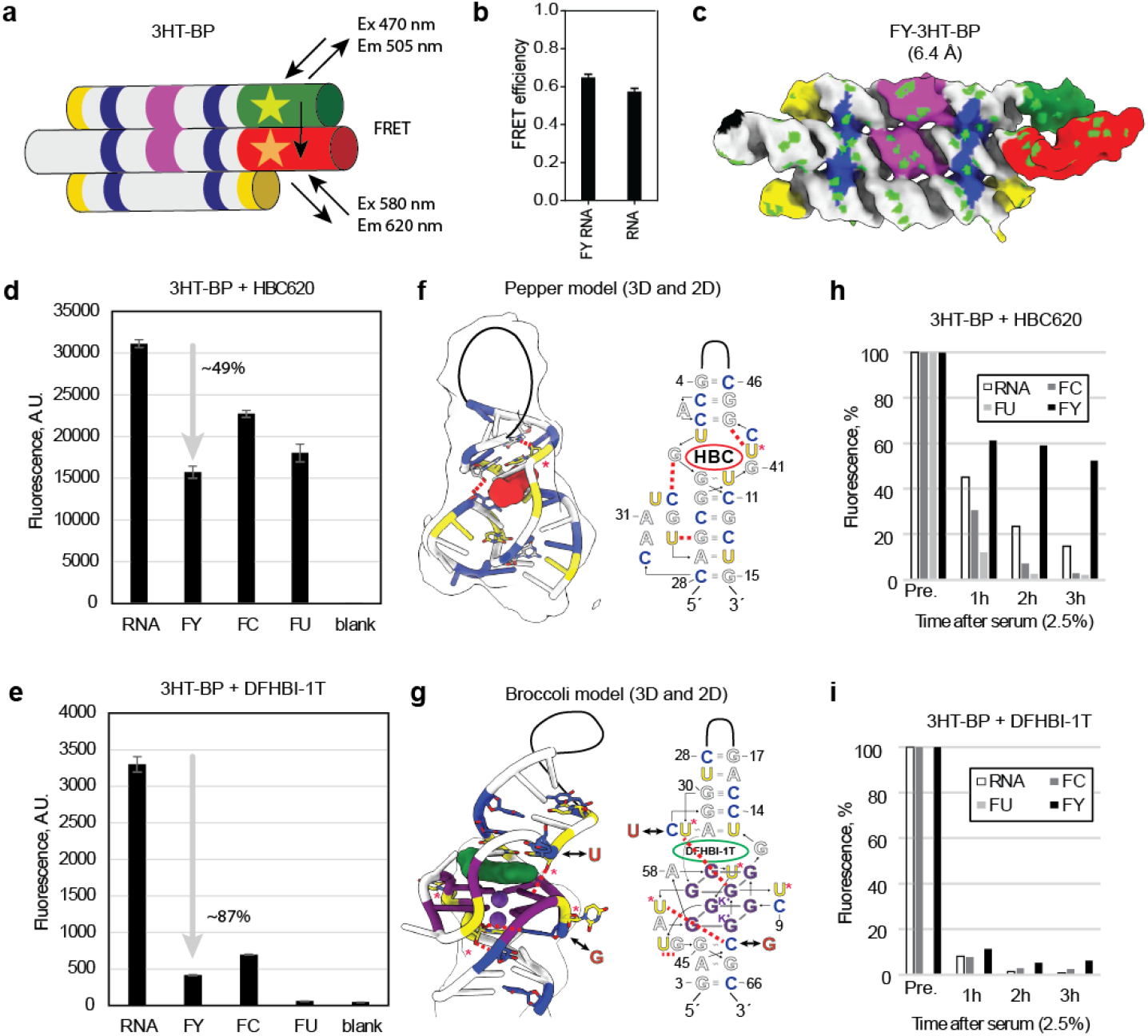
Fluorescence measurements of FY-RNA fluorogenic aptamers. **a**, Schematic of 3HT-BP showing excitation and emission wavelengths of Broccoli and Pepper and FRET. Motifs are coloured as in Fig. 1h. **b**, Quantification of FRET in FY-RNA/RNA 3HT-BP from fluorescence lifetime measurements of samples with either DFHBI-1T or both DFHBI-1T and HBC620 present. The data is presented as the mean ± standard deviation (n=3). **c**, Cryo-EM density map of the FY-RNA 3HB-BP origami resolved to 6.4 Å. Motifs are coloured as in Fig. 1h. Green spots depict the 2’-F. **d**,**e**, Fluorescence measurements of the 3HT-BP measuring Pepper fluorescence in the presence of HBC620 dye and Broccoli fluorescence in the presence of DFHBI-1T dye. **f**,**g**, 3D and 2D models of Pepper (PDB: 7EOM^60^) and Broccoli (PDB: 8K7W^61^) with C (blue) and U (yellow). Red dashed lines mark disrupted H-bonds. Red * marks possibly disrupted C2’-endo conformations. Numbering corresponds to crystal structures. The cryo-EM density is shown to indicate the region covered. **h**,**i**, Serum stability of Pepper and Broccoli as RNA, FC-RNA, FU-RNA and F-RNA.

To investigate Pepper and Broccoli individually, we transcribed the 3HT-BP as RNA, FU-RNA, FC-RNA and FY-RNA and measured fluorescence intensity when adding only one fluorophore at a time (Fig. 4d,e). The fluorescence of Pepper decreases by ∼49% as FY-RNA and to a lesser extent as FU- and FC-RNA (Fig. 4d) indicating an additive effect of FU and FC modifications on the fluorescence decrease. The fluorescence of Broccoli decreases by ∼87% as FY-RNA, to a lesser degree as FC-RNA, and completely as FU-RNA (Fig. 4e) surprisingly showing that FY-RNA rescues the disruptive effect of FU-RNA. The most likely explanation is that FU-RNA misfolds Broccoli, and FY-RNA restores this misfold because of the more even contribution to base pair stability when FC is present as well. Next, fluorescence lifetime measurements were used to investigate the molecular environment of the fluorophores^59^. For both Pepper and Broccoli minimal changes in fluorophore lifetime were observed when changing from RNA to FY-RNA (Fig. E7b, Table S6) indicating that the binding environments for the fluorophores are minimally affected. Together, this indicates that the FY-RNA versions of Pepper and Broccoli are destabilized compared to RNA but that a subpopulation can still fold to form functional binding sites for the fluorophores.

The fluorescence experiments were evaluated in the context of the known crystal structures of Pepper (PDB: 7EOM^60^) and Broccoli (PDB: 8K7W^61^). Structural analysis of Pepper shows that 2’-FY modification will disrupt three H-bonds: C33···G9, U42···G44, and U35···G37 (Fig. 4f, red dashed lines) and possibly affect the C2’-endo sugar pucker of U42 (Fig. 4f, red *). These structural changes explain the fluorescence data as an additive effect, where modification of FC-RNA disrupts one H-bond, FU-RNA disrupts two H-bonds, and FY-RNA disrupts three H-bonds fitting the observed decrease in fluorescence (Fig. 4d). Structural analysis of Broccoli shows that 2’-FY modification will disrupt three H-bonds: U31···G51, U36···C49, and U43···G44 (Fig. 4g, red dashed lines) and possibly affect the C2’-endo conformation of four U residues: U24, U50, U55, and U60 (Fig. 4g, red *). These structural changes explain the absence of fluorescence for FU-RNA, since several U 2’-OH interactions and conformations are affected. It also explains why FC-RNA Broccoli has higher fluorescence than FU-RNA, since no C 2’-OH interactions are lost. The high sensitivity of Broccoli to 2’-F modifications is likely the result of a highly constrained tertiary structure which is sensitive to small changes.

Finally, Pepper and Broccoli were tested for their function in human serum. When incubated in 2.5% human serum at 37 °C only the fluorescence of FY-RNA 3HT-BP is maintained over the 3-hour duration of the experiment (Fig. 4h,i). It is observed that the addition of serum decreases the fluorescent signal of FY-RNA Pepper to 60% and only decreases to 55% over 3 hours, whereas RNA, FC-RNA and FU-RNA gradually decrease to less than 15%. For Broccoli the FY-RNA signal is decreased to 10% in serum and drops to 5% over 3 hours, whereas RNA and FC-RNA decrease to undetectable levels, and FU-RNA is non-functional. Together, the experiments show that 2’-F modification of fluorogenic aptamers has negative effects on the tertiary structure and thus the fluorophore binding and fluorescence. On the other hand, the fluorogenic aptamers still function and the 2’-F modification stabilizes them against degradation by RNases in human serum.

### FY-RNA robotic device allows sensing in serum

Encouraged by the partial function of the FY-RNA fluorogenic aptamers, we tested the incorporation of 2’-F modifications into an RNA robotic device^46^, that enables reversible switching of the fluorogenic aptamer iSpinach^62^. The device is named “Traptamer”^46^ since it traps an aptamer in an RNA origami frame and can be activated by the addition of two RNA key strands, thus functioning as an AND gate (Fig. 5a). The overall structure of the FY-RNA Traptamer was validated by cryo-EM of the closed conformation at a resolution of 9.8 Å (Fig. 5b, Fig. S12). The function of the FY-RNA Traptamer was tested by the addition of RNA key strands, which resulted in increased fluorescence output and validation of the AND-gate function (Fig. 5c). This shows the ability of RNA key strands to invade the FY-RNA bKL via toehold-mediated strand-displacement despite the reported increased base pair strength of FY-RNA^31,34^. Importantly, the FY-RNA Traptamer has less leakage (fluorescence upon addition of only one key) and thus the AND-gate function is slightly improved compared to RNA (Fig. E8a,b). The improvement is most likely caused by the stronger base pairing reported for FY-RNA^31,34^ that will strengthen the 6 base pairs in the KL interaction keeping the Traptamer closed even when one KL has been “unlocked” by one of the RNA key strands. The decrease in leakage of the FY-RNA Traptamer fits with the improvement in binding strength of bKL-A and bKL-B (Fig. S6)^46^.

**Figure 5.**
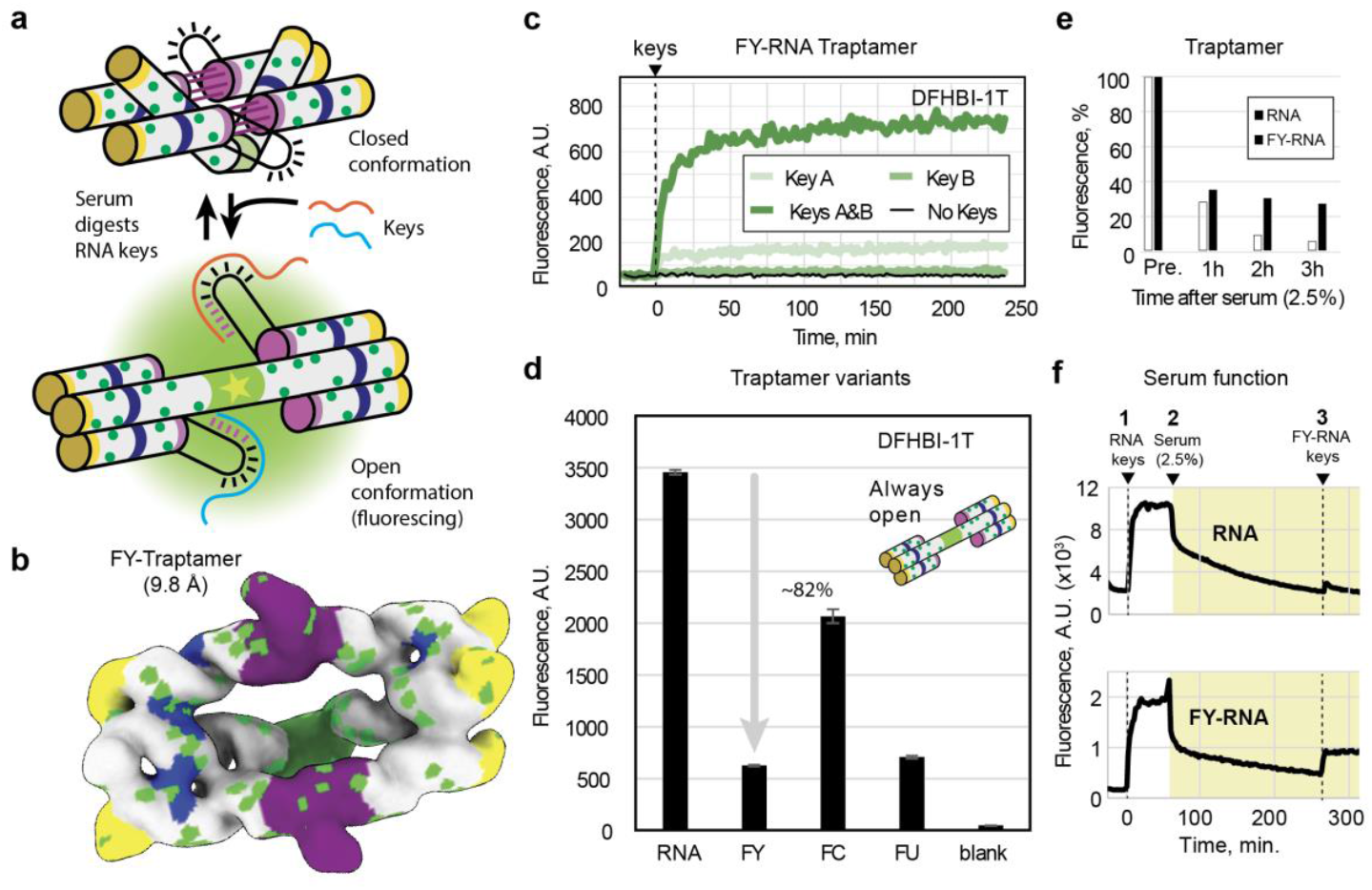
FY-RNA Traptamer stability and function in human serum. **a**, Illustration of the Traptamer in the closed conformation and how it can be opened by two RNA key strands to release and allow the iSpinach aptamer to binds its fluorophore and fluoresce. The Traptamer can be closed again by serum degradation of the RNA keys. **b**, Cryo-EM structure for the FY-RNA Traptamer at 9.8 Å resolution. **c**, Time-course experiment measuring fluorescence of the FY-RNA Traptamer when adding the keys individually or together in the presence of DFHBI-1T. **d**,**e**, Fluorescence measurements of an open-state variant of the Traptamer measuring iSpinach fluorescence in the presence of DFHBI-1T dye. **d**, iSpinach + DFHBI-1T in the Traptamer. **e**, Bar chart of normalized fluorescence of the RNA and FY-RNA Traptamer at the noted timepoints after adding 2.5% human serum. **f**, Time-course experiment showing the function of the RNA and FY-RNA Traptamer in serum when adding RNA keys at time zero, 2.5% human serum at 1 hour, and FY-RNA keys at 4.5 hours.

Fluorescence measurements of an open-state variant of the Traptamer, where iSpinach is functional, shows that FY-RNA decreases the fluorescence of iSpinach by 82% compared to the RNA version (Fig. 5d). Sequence comparison shows that iSpinach has two FCs less than Broccoli (C5G and C31U, Fig. 4g), which explains the 3x improved fluorescence of FC-RNA iSpinach compared to Broccoli (compare Fig. 4e and 5d). iSpinach has two additional FUs than Broccoli (C31U and A2U, Fig. 4g) but surprisingly FU-RNA iSpinach still fluoresces whereas FU-RNA Broccoli does not. Structural analysis of iSpinach (PDB: 5OB3^62^) shows that U31 is placed in a bulge and will likely not affect the structure, whereas G5 forms an important G5-G45 noncanonical base pair that is involved in stabilizing an additional potassium ion bound in a G-quartet^62^. This suggests, that G5 has a stabilizing effect on the FY-RNA iSpinach structure, that counteracts the negative effect of seven FUs, and increases fluorescence.^70^

Incubating the open-state Traptamer in 2.5% human serum at 37 °C results in reduced but persistent fluorescence for FY-RNA (Fig. 5e). Compared to Broccoli, the iSpinach in the Traptamer maintains 3-fold higher fluorescence in serum for 3 hours (compare Fig. 4i and Fig. 5e). Testing the Traptamer function in serum by sequential addition of RNA keys, 2.5 % human serum, and FY-RNA keys, revealed a turn-on, turn-off, and (only for the FY-RNA Traptamer) turn-on again. This indicates that the FY-RNA Traptamer functions in human serum, whereas the RNA Traptamer and RNA keys are degraded (Fig. 5f). The improved stability of the RNA sensor device was also tested in other challenging environmental samples such as lake water (Fig. E8c,d). In conclusion, it was shown that both Pepper, Broccoli, and iSpinach emit specific fluorescence after conversion to FY-RNA albeit at reduced intensity. In the future, FY-RNA fluorogenic aptamers can be improved by rational design or *in vitro* selection approaches^38^ in challenging environments such as active serum or waste water.

### Structure of FY-RNA aptamer binding the SARS-CoV-2 Spike protein

An FY-RNA aptamer has previously been demonstrated to bind and neutralize the SARS-CoV-2 virus through specific interaction with the receptor binding domain (RBD) of the Spike protein^14^. However, both the structure of the aptamer and the details of its interaction with the protein remained unknown. To solve the structure, we designed a FY-RNA PXT with the FY-RNA anti-Spike aptamer inserted at TL3 to make a protein-binding FY-RNA device (PXT-Spike) with a recognizable origami structure to function as a molecular ‘pointer’. Cryo-EM of the FY-RNA PXT-Spike incubated with the purified wildtype Spike protein resolved to 2.9 Å (locally to 2 Å) and reveals binding of the pointer device to three RBDs of the Spike protein trimer, which was found to be in its fully open conformation (Fig. 6a, Fig. S13-14). By C3 symmetry expansion and local refinement, the FY-RNA anti-Spike aptamer binding to the RBD was resolved to 3.4 Å revealing key molecular details such as individual bases and amino acid side chains (Fig. 6b). The FY-RNA PXT ‘pointer’ aided in building the anti-Spike aptamer atomic model (PDB: 9T74) by facilitating the positioning of the origami scaffold into the density map (Fig. 6c,d, Table S5).

**Figure 6.**
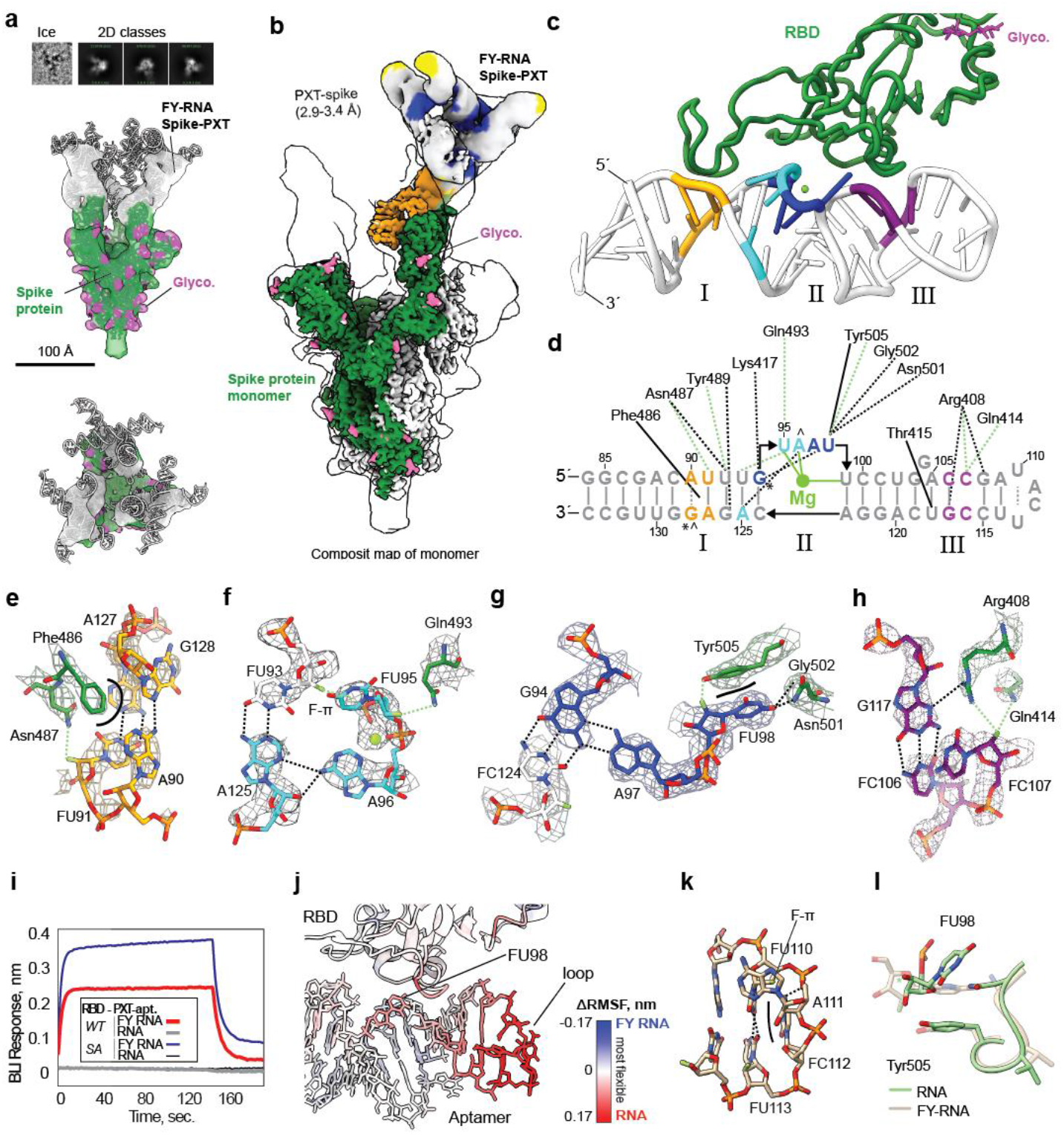
Structural analysis of FY-RNA anti-Spike aptamer using an origami ‘pointer’. **a**, examples of raw data particle, 2D classes and the full density map gaussian filtered to ∼10 Å for the trimeric Spike protein (green) with glycosylations (purple) and additional density in the top indicating binding of the FY-RNA origami pointer (white). **b**, Composite map of the best resolved maps from reconstruction of the symmetry expanded Spike protein monomer. Transparent density represents the best full non-symmetry enhanced map filtered to a resolution of ∼10 Å. **c**,**d**, Tertiary and secondary structure of the FY-RNA anti-Spike aptamer interacting with the RBD of the Spike protein. * marks C2’-endo sugar pucker. ^ marks syn conformation. Black dashed lines indicate H-bonds. Green dashed lines indicate 2’-F specific interactions. Black lines indicate hydrophobic interactions. Mg^2+^ ion shown as green circle. **e**-**h**, Zoom-in on binding regions I-III shown in the cryo-EM focus map. Colours correlate with those in panel c,d. **i**, Bio-layer interferometry measurement of the binding between immobilized RBD of wild type (WT) and South African variant (SA) SARS-CoV-2 Spike proteins with the FY-RNA PXT-Spike aptamer. **j**, MD simulation data plotted as ΔRMSF on the aptamer structure comparing RNA with FY-RNA. **k**, MD simulation shows stabilization of AUCU TL in FY-RNA but not in RNA. **l**, MD simulation show interaction between U98 and Tyr505 in FY-RNA (brown) but not in RNA (green).

From the cryo-EM structure it is clear that the binding of the anti-Spike aptamer blocks the binding of the RBD to the ACE2 receptor. The anti-Spike aptamer binds to the RBD with three regions, termed region I, II, and III (Fig. 6c,d). Region I has a cis-Watson-Crick-Hoogsteen base pair between A90-G128 with the G base in syn conformation, which creates a hydrophobic pocket in the shallow groove for interaction with Phe486 (Fig. 6e, black line). A stabilizing electrostatic interaction is found between the 2’-F of FU91 and the side chain amine of Asn487 at a distance of 3.7 Å (Fig. 6e, green dashed line). Tyr489 contacts a G-U base pair in the shallow groove, where it can H-bond with G126 and has an electrostatic interaction with the 2’-F of U92 at a distance of 3.8 Å (not shown). Region II has a 4-base bulge that projects out of the helix after G94 (Fig. 6d), which is arranged around a centrally bound Mg^2+^ ion that coordinates with the phosphodiester backbone of FU95, A96, and likely FU99 and results in the outward projection of FU95 and FU98. The bulge is further stabilized by two A-minor interactions of A96 and A97 (Fig. 6f,g). Lys417 H-bonds to the backbone of G94 (not shown). The motif promotes specific interactions with the protein by forming a hydrophobic interaction between FU95 and Phe456 (Fig. 6f), which may be further stabilized by the 2’-F of FU93 that points towards the base of FU95 and is in position to make a F-π interaction at ∼4 Å distance^63^ (Fig. 6f, F-π). A stabilizing electrostatic interaction is found between the 2’-F of FU95 and the side chain amine of Gln493 at a distance of 3.6 Å (Fig. 6f, green dashed line). The outward projecting FU98 has a stacking interaction with Tyr505 and makes two H-bonds with the backbone and sidechain amine of Gly502 and Asn501, respectively (Fig. 6g). The 2.6 Å distance between 2’-F of FU98 and the hydroxyl group on Tyr505 likely represents a genuine F-based H-bond that is stabilized by the stacking interaction (Fig. 6g). Region III contains a C107-C116 mismatch that allows for a shallow groove interaction with the side chain amines of Arg408 that makes H-bonds to FC107, G108, and G117 (Fig. 6h). Further, stabilizing electrostatic interactions at ∼3.5 Å are found between the 2’-F of FC107 and side chain amines of Arg408 and Gln414 (Fig. 6h, green dashed lines). 2’-F of FU118 makes a hydrophobic interaction with the methyl side chain of Thr415 with a distance of 3.0 Å. In summary, we find five electrostatic interactions of 2’-F with H-donors at ∼3.6 Å, a possible F-π interaction at ∼4 Å, a 2’-F hydrophobic interaction at 3.0 Å, and one F-based H-bond at 2.6 Å.

Biolayer interferometry (BLI) analysis shows that the FY-RNA PXT-Spike binds strongly to the Spike RBD of both the SARS-CoV-2 wildtype (WT) and South African (SA) variants, whereas the RNA version of the aptamer does not bind the Spike proteins at all (Fig. 6i). To investigate why, we performed MD simulations starting from the cryo-EM structure of the anti-Spike aptamer bound to the RBD simulated as both FY-RNA and RNA (Fig. 6j, Fig. E9, Fig. S21 and Video 4). The simulation identified the loop region UACU113 and U98 to be much more flexible as RNA compared to FY-RNA (Fig. 6j). Inspection of the loop region reveals that FU110 and FU113 form a cis-Watson-Crick-Watson-Crick base pair, C112 stacks on top, and A111 makes contact with the 2’-F of U110 forming a potential F-π interaction^63^ (Fig. 6k). The UACU sequence may represent a FY-RNA specific TL that can be used to improve future FY-RNA designs. In RNA the stacking interaction between U98 and Tyr505 was disrupted caused by U98 adopting a C2’-endo conformation (Fig. 6l, Fig. S21) and interaction with the backbone of U99 and C100 by the 2’-OH (Supplementary Videos 5-6). The stable interaction between FU98 and Tyr505, observed in the FY-RNA simulations and cryo-EM data, is likely caused in part by increased base stacking in 2’-F-modified residues^31,34^, the 2’-F H-bond interaction with the OH of Tyr505, and by the reduced reactivity of the 2’-F of U98 with the nearby backbone. It has previously been shown that the anti-Spike-aptamer does not bind the Omicron variant^14^. Interestingly, the Omicron variant has a mutation of Tyr505 to Histidine (T505H) (Fig. E5b,c) that may perturb the specific stacking and bonding interaction of FU98. This potentially explains the inability of the anti-Spike-aptamer to bind the Omicron Spike protein and highlight the importance of this 2’-F stabilized interaction. The atomic structure presented here can potentially inform rational redesign or guide *in vitro* selection strategies to generate next-generation Spike-binding aptamers capable of efficiently targeting Omicron or future viral variants.

## Conclusions

Based on our findings, we can conclude that FY-RNA folds similarly on the secondary structure level but that F-specific effects arise at the tertiary structure level. Similarities of secondary structures is likely the result of using modifications on both C and U, which provides an even stabilization of canonical base pairs resulting in a similar but more stable energy landscape. Differences seen for tertiary structures were found to be a consequence of how 2’-F modification increases stability of base pairing and stacking, disrupting H-bonds, favouring the C3’-endo sugar conformation, and increasing the hydrophobicity of the shallow groove. The 2’-F modification led to dramatic folding effects in the 6HB and to decrease in fluorescence of fluorogenic aptamers. On the other hand, when determining the structure of an anti-Spike aptamer that was originally selected as FY-RNA, we found FY-RNA tertiary motifs and protein interactions that are unique and takes advantage of the properties of the F atom. Interestingly, we find that 2’-F often engages in long-range electrostatic interactions and sometimes as an acceptor for H-bonds. Together, the study allowed us to derive several design principles for FY-RNA origami and expanded the toolbox of FY-RNA modules that can be used for construction of FY-RNA origami devices. In the future, more FY-RNA selections are needed to either convert existing RNA motifs into functional FY-RNA versions or to select FY-RNA functionality such as aptamers, riboswitches and ribozymes.

Our characterization platform is generalizable to the study of other XNAs and can be used to explore the structure and functional landscape of enzymatically produced XNA polymers. As an example the structural effects of base-modifications of the 6HB has been recently demonstrated^52^. The 6HB and PXT-Spike-aptamer structures of this study were submitted to the Critical Assessment of Structure Prediction (CASP16) competition, where it was found that 2’-F is currently not supported and our structures were not correctly predicted^65^. This highlights that more XNA nanostructures should be structurally determined to provide sufficient information for training machine-learning algorithms to predict and design these polymers - and for us as rational designers to understand the core design principles. To expand our characterization platform to new XNA nanostructures the following is needed: (1) Engineered polymerases able to efficiently synthesize XNA. (2) Dedicated design software that incorporates the energy parameters of XNAs to allow secondary structure prediction, which is currently being developed for modified nucleotides for the ViennaRNA Package^66^. (3) Methods for structure determination and precise modelling, where our NAMiX software facilitates model conversion from RNA to XNA. (4) Force fields for XNAs for MD simulation. Together, these developments will help provide a fundamental understanding on the role of chemical modifications, both in biology, biotechnology, and therapeutics.

## Material and methods

### Design and production of FY-RNA origami structures

The construct-design is done within the frames of the standard RNA origami design strategy, which was previously described for RNA^44,50^ but adapted for our emerging understanding of FY-RNA. Briefly, a sequence generalized 2D blueprint describing the desired base pairing, modular composition and strand path including KL contacts but not the specific base pair identity is input into the Revolver program from the ROAD software suite^50^ generating many sequences that fulfil the requirements. Based on these, the sequences with the lowest ensemble diversity (as if they were RNA)^50^ are identified and manually checked in NUPACK^67^ (as if they are RNA).

The best candidate sequence is then ordered as double-stranded DNA for direct transcription or for cloning into plasmids and transcription from the plasmid. DNA is sequenced to ensure the correct sequence is present. For large scale transcription, large amount of plasmid is produced in DH5-aplha cells and extracted using a maxiprep kit (Macherey-Nagel). Purified plasmid is then restriction digested overnight cleaving the DNA so that the last template nucleotide is the 3’-end of the origami. Then the linearized plasmid DNA is purified by phenol-chloroform extraction followed by ethanol precipitation and dilution in water. Alternatively, the double-stranded DNA is directly amplified by PCR using Q5 DNA polymerase (New England Biolabs) followed by PCR clean up (Qiagen).

Transcription reactions are set up in a transcription buffer containing 40 mM HEPES (pH 8.0), 25 mM MgCl_2_, 2 mM Spermidine 50 mM KCl, 2.5 mM GTP, 2.5 mM ATP, 2.5 mM 2’-F-UTP, 2.5 mM 2’-F-CTP, 12.5 mM DTT, 0.05mg/mL BSA, Ippase, and 100 nM DNA template. Transcription is then started by addition of mutant (Y639F) T7 RNA polymerase^37^ prepared in-house or purchased (Aptamist ApS). Transcription is carried out for 3-16 hours at 37 ^°^C. Transcription reaction is then loaded on to a Superose 6 column (Cytiva) equilibrated with 40 mM HEPES buffer (pH 8), 50 mM KCl and 5 mM MgCl_2_. The fractions containing the purified sample (evaluated by the approximate expected elution time based on the elution time of the other origamis of varying size) is then collected and concentrated in a 30 kDa-cutoff spin filter (Amicon) to a concentration between 1-50 µM and analysed right away or frozen.

### Negative stain-EM data acquisition

For a negative stain electron microscopy (ns-EM), the purified and concentrated samples were diluted 10-fold. We use locally prepared grids (400 mesh copper grids with collodium and carbon) that were glow discharged for 45 s at 25 mA using a PELCO easiGlow before sample application. Then 3 µL of the sample was applied to the carbon film for 30 sec before side blotting on Whatman #1 filter paper followed by immediate application of 3 µl of uranyl formate (5%) solution, blotting after 15 sec, and reapplication of uranyl formate for a total of three rounds of staining. The grids were then air-dried for 10 min before imaging. The images were acquired on Tecnai Spirit at 120 kV equipped with a TVIPS 4k camera with a pixel size of 1.54 Å px^−1^. The data were saved as 16-bit TIFF files and manually inspected.

### Cryo-EM data acquisition

Purified and concentrated samples were taken immediately for grid plunging. We used Protochips 1.2/1.3 300 mesh Au-Flat grids (Jena bioscience) that are glow discharged in a GloQube Plus glow discharging system for 45 seconds at 15 mA and used within ∼30 min for plunge freezing. Plunge freezing was performed on a Leica GP2 with the sample chamber at 90% humidity and 5 °C. 3 µL samples were applied onto the foil side of the grid in the sample chamber before a 4 second delay and then 6 seconds of distance-calibrated foil-side blotting against a double layer of Whatman #1 filter paper. With no delay after blotting, the sample was plunged into liquid ethane set to −184 °C and kept under liquid nitrogen until clipping and loading into the microscope. All data were acquired at 300 keV on a Titan Krios G3i (Thermo Fisher Scientific) equipped with a K3 camera (Gatan/Ametek) and energy filter operated in EFTEM mode using a slit width of 20 eV. Data were collected over a defocus range of −0.8 to −2 micrometers with a targeted dose of 60 electrons per square Ångstrom (Å). Automated data collection was performed with EPU, and the data was saved as gain normalized compressed tiff files with a calibrated pixel size of 0.647 Å per pixel.

### Single-particle image processing and 3D reconstruction of cryo-EM data

The recorded movies were pre-processed with cryoSPARC which performed motion correction, contrast transfer function (CTF) correction as well as the micrograph curation and downstream analysis for particle picking, and structural analysis. For all samples the data was f-cropped to ½ before motion correction (1.294 Å per pixel) and only good micrographs were selected in the manual curation to be used in blob picking which together with 2D classification and *ab-initio* reconstruction was used to generate the initial templates for template picking. Template picked particles were analysed in detail to generate refined density maps. For datasets with potential for good resolution, Topaz picking was trained on a subset of the particles (and micrographs) from a refined density map and used to extract particles form the whole dataset. These particles were analysed extensively to reach the best possible stack for reconstruction. For symmetric particles we ran parallel analysis both with or without the relevant symmetry imposed, followed by symmetry expansion, particle subtraction, and local refinement of the focus areas. 3D classification, 3D variation, 3D flex analysis tools were explored to access the structural classes and variation in the particle stacks. Some of the maps reached the Nyquist limit of the cropped micrographs (∼2.5 Å). In those cases, we re-extracted particles from uncropped motion correction and CTF fitted micrographs followed by re-analysis reaching resolution down to ∼2 Å (Nyquist is ∼1.3 Å). Relevant data, maps, and models are available on EMPAIR, EMDB and PDB.

### XNA models with NAMiX

The presented FY-RNA origami models were generated based on the cryo-EM density maps with knowledge of the designed structure and base pairing pattern of the origami as well. The density map resolution varied from 10-2 Å (GSFSC, 0.143). Between 10 and 4 Å resolution, the maps were used to trace the strand path and confirm the overall folding and organization of the origami. Detailed investigation of these models relies on MD simulation. For resolutions below 4 Å, models are evaluated in greater detail describing specific molecular interactions if supported in the data.

Initial models of RNA were derived either from existing RNA origamis, made using RNAbuild in the ROAD software suite^50^, or build from scratch using UCSF ChimeraX (version 1.9)^68-70^. The models (still RNA) were then fitted into the map densities in UCSF ChimeraX using ISOLDE (version 1.9) to perform MD with flexible fitting with a low (500 kJ mol^−1^ (map units)^−1^ Å^3^) weight on the map and 0 as the temperature factor^71^. For maps with resolutions between 10-4 Å, it is very helpful to initially make sure to use distance restrains to specify base paring (3 Å, spring constant: 50 kJ mol^−1^ Å^−2^), base stacking (4 Å between bases, spring constant: 50 kJ mol^−1^ Å^−2^) and phosphate distances (5.9Å, spring constant: 50 kJ mol^−1^ Å^−2^) before starting the simulation.

After ISOLDE, the models were then converted to FY-RNA using the (made for the purpose) NAMiX software package, which also generates blueprint-based secondary structure restrains and outputs eff and cif files for further model refinement. The FY-RNA models were then real-space refined (<4 Å) or geometry minimized (≥4 Å) using the Phenix software package^72^ and eff and cif files from NAMiX to optimize bond angles and reduce the clash score before final validation using Phenix. Validations of the models are presented in Table S2-5. NAMiX was also developed for locked-nucleic acid (LNA) and can later be extended with any modified nucleotide.

### System preparations for MD-simulation

System building was initiated from cryo-EM informed structures of the PXT and 5HT and from structural motifs in these: TL and DX of the PXT and KL of the 5HT. Chemical modifications were introduced into RNA versions of the structures using NAMiX. tleap was used to add hydrogens and create topology files for simulations in the AMBER force field (see Amber 2023 reference manual, version 2023). The Python package acpype was used to convert AMBER force field files into GROMACS format^73^. We used GROMACS version 2024.3 for both system building and MD simulations (see GROMACS 2024.6 Manual, version 2024.3, zenodo). The systems were solvated in dodecahedral boxes with a solvation distance of 1.2 nm using gmx solvate. Ions were added and the system was neutralized with 50 mM MgCl_2_ using gmx genion. The systems were energy minimized using the steepest descent algorithm for 10,000 steps. Two rounds of NPT equilibrations were performed using the Nosé-Hoover thermostat at 310 K and with position restraints of 1000 kJ mol^−1^ nm^−2^. The first equilibration step was performed using a timestep of 1 fs and 125,000 steps yielding 0.125 ns, and the second was performed using a 2 fs timestep and 25,000,000 steps resulting in a 50 ns equilibration. The system setup can be reproduced by using the method described in https://github.com/AmandaStange/RNA.

### MD simulations

The MD simulations were performed using the AMBER force field and specifically ff19SB was used for proteins^74^, parmbsc1 for RNA^74^, and f-sugars for fluor-modified sugars^47^. The PME algorithm was used for electrostatic interactions with a cut-off of 0.9 nm^75^. A single cut-off of 0.938 nm was used for Van der Waals interactions. The pressure was maintained at 1 bar using the Parinello-Rahman barostat^76^ and the temperature was kept at 310 K using the v-rescale thermostat^77^. Four repeats were simulated with different starting velocities for 500 ns each using a timestep of 2 fs.

### MD simulation analysis and statistics

The RNA and protein RMSD along the trajectories were calculated using gmx rms and gmx rmsf was used to calculate their RMSF (GROMACS 2024.3 Manual, version 2024.3, zenodo). ProLIF was used to calculate the hydrogen bond information along the trajectories^78^. The python package biobb_dna was used to calculate the helical parameters (rise, twist, inclination) and sugar parameters (puckering)^79^ over the trajectories. Crossover angles were calculated using MDTraj^80^ and PCA from scikit-learn^81^. Heavy atoms of each helix were defined as point clouds and PCA was used to fit a vector describing the direction of the helices. The angle between the PCA vectors was calculated hereby yielding the crossover angle over the trajectories. Trajectory frames from the last 200 ns were clustered using all-against-all heavy-atom RMSD and average-linkage hierarchical clustering implemented in SciPy^82^, with the number of clusters selected by an elbow criterion refined by silhouette analysis using scikit-learn^81^; representative structures were defined as cluster medoids with minimal mean intra-cluster RMSD. The gaussian_kde method from SciPy^82^ was used to generate the angle distributions. Data sets presented as means stem from meaning over the four repeats and the error bars represent the standard deviation, both calculated using the python package NumPy^83^. Plotting was done using the Python packages matplotlib^84^ and Seaborn^85^, molecular visualization and mapping was performed using ChimeraX^68-70^.

### Gel based serum stability assay

Purified FY-RNA origamis or the corresponding RNA origamis were incubated at a final concentration of 0.93 µM in a mixture containing 25% human serum, 25% Dulbecco’s Phosphate Buffered Saline (DPBS), and 50% origami buffer (40 mM HEPES, 50 mM KCl, and 5 mM MgCl_2_) at 37 °C. Aliquots (0.5 µL) were collected at various time points (1 min, and 3, 6, 24, 48, 72 and 144 hours), diluted in 4.5 µL MilliQ water, and 2.5 µL (corresponding to 0.625 pmol) were analysed by gel electrophoresis on an 8% denaturing polyacrylamide gel. Gels were quantified using ImageJ software (https://imagej.nih.gov/ij/).

### RNase A digestion assay

Purified FY-RNA and RNA origamis were incubated in 40 mM HEPES, 50 mM KCl and 5 mM MgCl_2_ at a final concentration of 0.5 µM with or without 1 ul RNase A (20 mg/mL, NEB) for 1 hour at 37 °C. The results were analyzed by gel electrophoresis on an 8% denaturing urea polyacrylamide gel.

### Fluorescence detection

Measurements were performed in 384-well plates (Costar) with sample volumes of 40 μl containing 25 nM sample (RNA, FU-RNA, FC-RNA, and FY-RNA), 500 nM DFHBI-1T (Lucerna Technologies) and / or HBC620 (FR Biotechnology), 40 mM HEPES, 50 mM KCl, and 5 mM MgCl_2_. Fluorescence was monitored using a CLARIOstar Plus multi-mode microplate reader (BMG LABTECH) set at a constant temperature of 37 °C. For the addition of human serum, RNA keys or FY-RNA keys, the plates were quickly removed from the plate reader, adding and mixing the new components (5 µL each time), and then the plates were reinserted into the machine for continued measurements. Addition of these components slightly changed volume of the system as well as the buffer and component concentrations. Importantly, fluorescence is measured in the whole volume, thus the fluorescence output is not varying due to dilution. Excitation of DFHBI-1T was performed at 470 ± 10 nm and emission was recorded at 505 ± 10 nm for the time point measurements. Excitation of HBC620 was performed at 580 ± 10 nm and emission was recorded at 620 ± 10 nm for the time point measurements.

### Fluorescence lifetime measurements

Fluorescence lifetime measurements were recorded in-solution using a Luminosa single photon counting confocal microscope (PicoQuant). FY-RNA samples contained 500 nM FY-RNA, 2.5 μM DFHBI-1T and/or 2.5 μM HBC620. RNA samples contained 200 nM RNA, 1 μM DFHBI-1T and/or 1 μM HBC620. A standard buffer of 40 mM HEPES, 50 mM KCl, 5 mM MgCl_2_, and 0.01 % Tween20 was used for all measurements. For each measurement, a 30 uL droplet was placed on a coverslip. DFHBI-1T was excited with a 488 nm laser (PicoQuant, LDH-D-C-485S), and its emission was collected with a 511/20 band pass filter. HBC620 was excited with a 530 nm laser (PicoQuant, LDH-D-FA-530L), and its emission was collected with a 690/70 band pass filter. The laser power was approximately 4 μW and the repetition rate was 16 MHz. Due to the low fluorescent signal from FY-RNA, FY-RNA measurements were recorded for 10 min, whereas measurements with RNA were recorded for 2 min. All measurements were repeated 3 times. The fluorescence decays were analysed in the Luminosa Analysis software (PicoQuant), where n-exponential reconvolution was used to extract the lifetime components and their relative amplitudes.

### Fluorescence spectra measurements

Fluorescence spectra measurements for assessing FRET between the Broccoli and Pepper aptamers were conducted on a FluoroMax-3 fluorometer (Jobin Yvon, Horiba). Samples contained 200 nM FY-RNA, 4 μM DFHBI-1T, and/or 2 μM HBC620, 40 mM HEPES, 50 mM KCl, 5 mM MgCl_2_, and 0.01 % Tween20. A 60 μL 3×3×10 mm quartz cuvette (Hellma Analytics) was used. A slit range of 5 nm and an integration time of 0.5 s were used to scan all spectra. Spectra were recorded in ratio mode that corrects for wavelength-dependent detection sensitivities by normalization of the measured signal against a reference detector signal. DFHBI-1T was excited at 460 nm and emission was collected in the spectral range 470-740 nm. HBC620 was excited at 550 nm and emission was collected in the spectral range 560-740 nm. Corresponding backgrounds of free fluorophore in 40 mM HEPES, 50 mM KCl, 5 mM MgCl_2_, and 0.01 % Tween20 were subtracted from all measurements.

### FRET analysis

FRET efficiencies, *E*, were determined by *E* = 1 − *τ*_*DA*_/*τ*_*D*_, where *τ*_*DA*_ is the amplitude weighted average DFHBI-1T lifetime measured in an RNA or FY-RNA sample containing both DFHBI-1T and HBC620, while *τ*_*D*_ is the DFHBI-1T lifetime measured in an RNA or FY-RNA sample containing only DFHBI-1T.

### BLI binding measurements

Biolayer Interferometry (BLI) experiments were performed using the Octet RED96 system (ForteBio, Sartorius). All measurements were conducted in black, flat-bottom 96 well plates (Grenier) with orbital shaking at 1000 rpm and at room temperature (RT). To evaluate the binding of PXT-Spike as FY-RNA and RNA, two His-tagged variants of the SARS-CoV-2 receptor binding domain (RBD) Spike protein (wild-type (WT) and B.1.351 (Beta, South Africa)) were diluted to a final concentration of 2.5 µg/mL in binding buffer (PBS pH 7.4, 3 mM MgCl_2_, supplemented with 0.1 mg/mL BSA and 0.02% Tween-20) and immobilized onto Ni-NTA–coated biosensors (OCTET Ni-NTA [NTA] Biosensors, Sartorius). PXT-Spike were tested at a concentration of 600 nM in binding buffer. Prior to each binding experiment, a baseline was recorded in buffer to establish initial BLI signals. Subsequently, the protein-coated sensors were dipped into the PXT-Spike solutions (association phase), followed by immersion in buffer-only wells (dissociation phase). Three cycles of regeneration were performed by dipping the sensors into 10 mM glycine solution (pH 1.4) followed by buffer, each for 5 sec. An additional surface recharging step was carried out by dipping the sensors in 10 mM NiCl_2_ in water for 30 sec. Sensorgrams were analysed using the Octet software.

## Acknowledgments

We thank the EMBION Cryo-EM Facility at iNANO, Aarhus University, for time on the Titan microscopes, technical support, and data processing on the EM computer cluster. Thanks to Rita Rosendahl and Claus Bus for technical assistance. We thank the Centre for Scientific Computing in Aarhus for access to the computational resources.

## Author contributions

ELK, CG, JK, JV and ESA designed research; ELK, NZ, NSV, NG, LC, CG, ADS, LMD and ST performed the research; ELK and NZ performed the structural biology; ADS and LMD performed the MD simulations; ST and VB performed the fluorescent life time and FRET measurements; all authors helped analyse the data; and ELK and ESA wrote the paper. All authors read and commented on the text.

## Data and materials availability

NAMiX version 1.0 available on Github: https://github.com/esa-lab/NAMIX. MD simulation scripts available at https://github.com/AmandaStange/RNA and MD simulation data at: https://zenodo.org/uploads/17938024. Cryo-EM data will be made available on EMPIAR (https://www.ebi.ac.uk/empiar/). Cryo-EM maps and models produced in this work are available on EMDB and PDB with the following IDs: PXT (EMD-53787, 9R7Q), 5HT (EMD-53795, 9R7W), 6HB (EMD-53803, 9R82), 6HB-dimer (EMD-54779, 9SD9), 3HT-BP (EMD-55626, NA), Traptamer (EMD-55635, NA), PXT-Spike (EMD-55622, EMD-55624, EMD-55621, EMD-55623), Anti-Spike-aptamer (EMD-55625, 9T74).

