## Supplementary materials for "Fluorinated RNA origami enables serum-stable nanodevices for sensing and targeting"

**Affiliations:**
[1]Interdisciplinary Nanoscience Center (iNANO), Aarhus University, 8000 Aarhus, Denmark.
[2]Department of Molecular Biology and Genetics, Aarhus University, 8000 Aarhus, Denmark.
[3]Department of Chemistry, Aarhus University, 8000 Aarhus, Denmark.

**Current affiliations:**
[4]Department of Health Technology, Danish Technical University, 2800 Lyngby, Denmark
[5]Department of Bioscience, School of Natural Sciences, Technical University of Munich, 85748 Garching, Germany.
[6]Center for Molecular Biology of Heidelberg University (ZMBH), University of Heidelberg, 69120 Heidelberg, Germany.

*Corresponding authors.

### Extended data figures

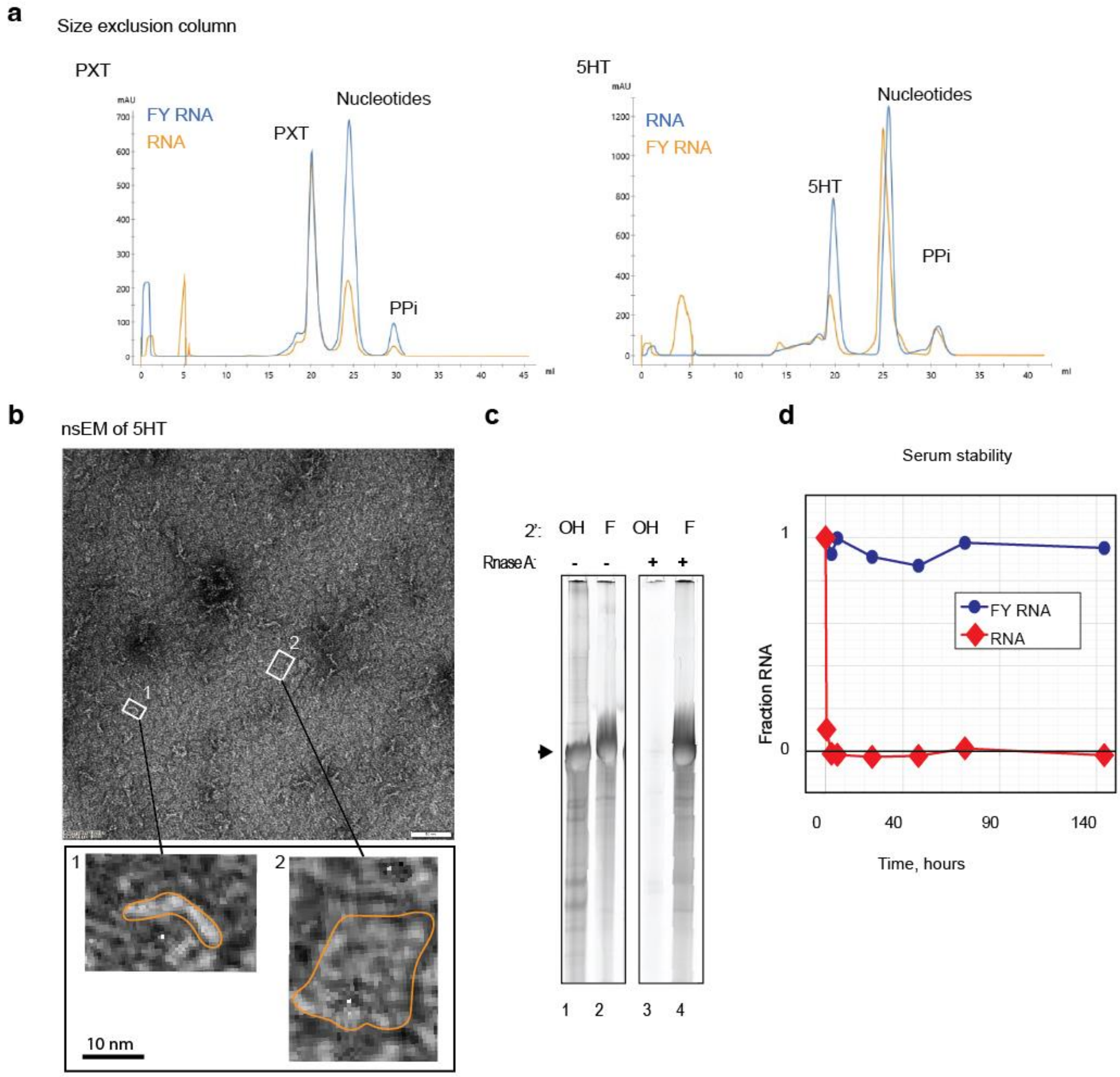

**Figure E1. Production, purification and initial analysis of FY-RNA structures. a**, Elusion curves from size exclusion purification of selected origami designs, PXT and 5HT, showing both FY-RNA and RNA. In both cases the peaks overlap indicating that the molecules are the same size and form well-defined particles (non-folding origamis often aggregate due to their many complementary regions and elude very differently). **b**, Image from negative stain electron microscopy (nsEM) analysis of the purified peak fraction from the FY-RNA 5HT size exclusion purification in panel a. The image shows a carpet of square-like structures resembling FY-RNA 5HT lying flat on the surface or standing on the edge (see zoom-in images 1 or 2). **c**, Denaturing PAGE analysis of RNase A treated RNA and FY-RNA origami samples. Lanes 1 and 2 represent untreaded samples, in lanes 3 and 4 samples were pre-incubated with RNase A. **d**, Quantification of denaturing PAGE analysis of origami samples pre-incubated with 50 % active human serum for the noted time. RNA was digested within the first 30 minutes; FY-RNA was still present after 140 hours.

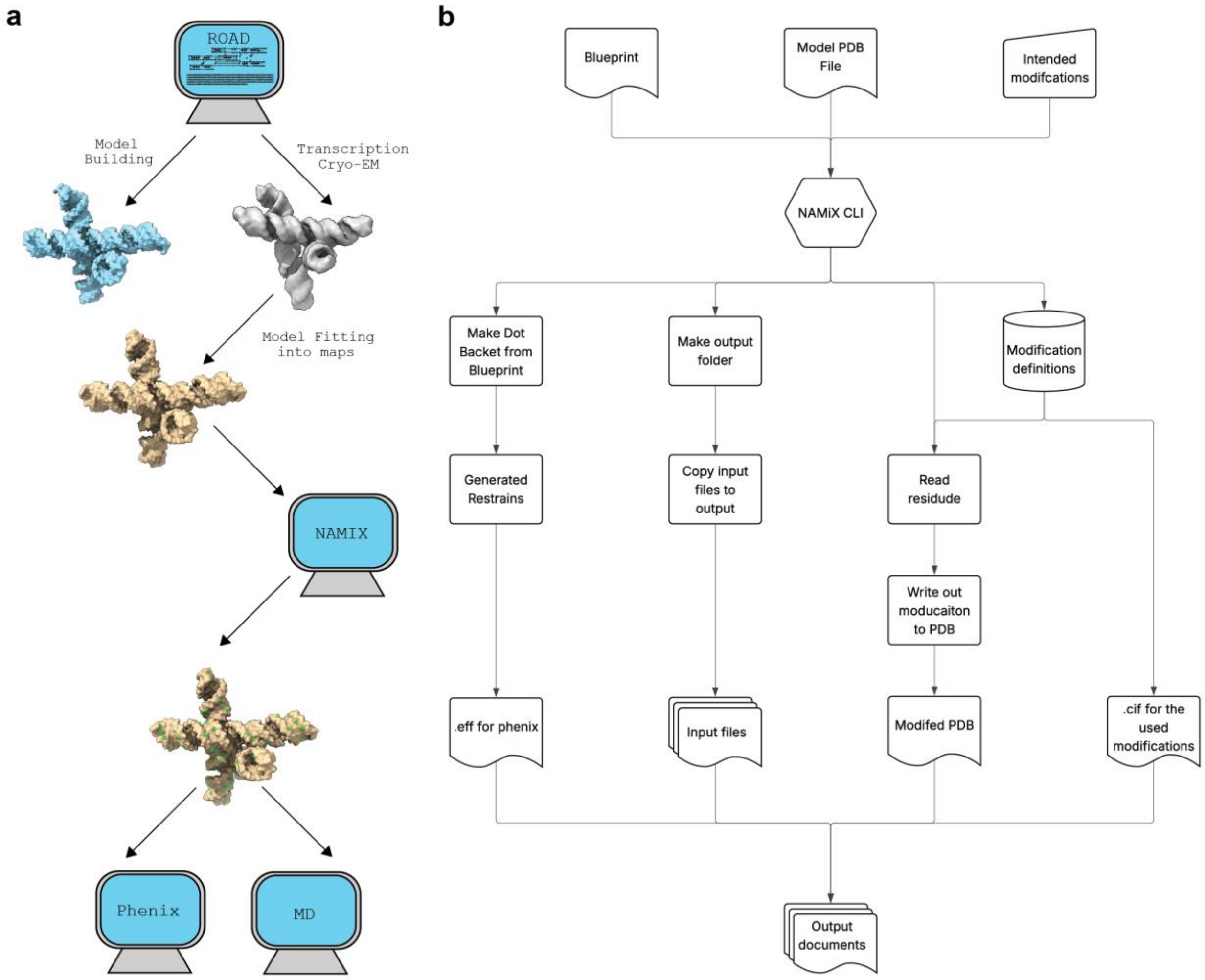

**Figure E2. NAMiX software. a**, Workflow for generation of Cryo-EM informed models. For the generation of Cryo-EM informed models, the workflow used in this paper utilizes the ROAD software package for sequence design and initial model generation followed by EM data acquisition. Then after initial fitting of RNA models into the maps, the models are processed into FY-RNA by NAMiX followed by validation and analysis with Phenix and molecular dynamics. **b**, Simplified data flow in the NAMiX software. Flowchart showing the overall data flow in the NAMiX software highlighting the different parts of the software. The primary of these being code for generation of modified PDB files. Second, there is the code for base pair and stacking generation code using ROAD-based blueprints. Lastly, is the "bookkeeping" code for transfer and organization of the used and generated files.

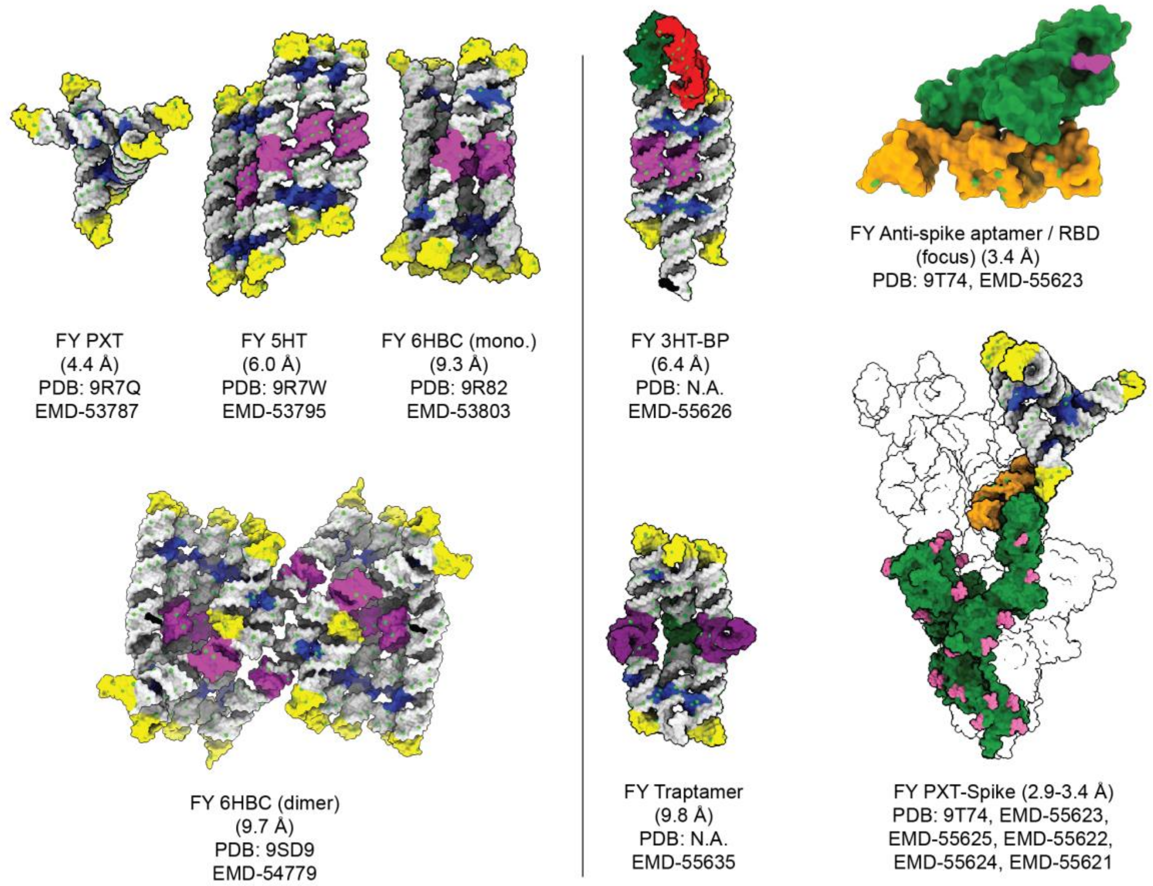

| Blueprint | FY-RNA PDB | RNA PDB | FY-RNA EMD | RNA EMD |
|---|---|---|---|---|
| PXT | 9R7Q | 8BTZ | 53787 | 16244 |
| 5HT-TC | 9R7W | 7QDU | 53795 | 13926 |
| 6HBC | 9SD9 (dimer) | 7PTL (mature)<br>7PTK (young) | 54779 (dimer) | 13625 (mature)<br>13626 (young) |
| 6HBC-TC | 9R82 (monomer) | ND | 53803 (monomer) | ND |
| 3HT-BP | ND | 7ZJ4 (bound)<br>7ZJ5 (unbound) | 55626 (bound) | 14740 (bound)<br>14741 (unbound) |
| Traptamer | ND | 8TVZ | 55635 | 41656 |
| PXT-Spike | 9T74 | ND | 55625 (RBD-aptamer)<br>55622 (Spike core)<br>55624 (NTD)<br>55621 (PXT)<br>55623 (Full map) | ND |

**Figure E3. Cryo-EM informed atomic models of the FY-RNA origamis and devices.** Models of the FY-RNA structures investigated with cryo-EM. Structural motifs (as shown in the main text figure 1G) are indicated by coloured domains. Position of the F-atoms is indicated by lime-green. In the spike protein (showing only one of the monomers), the amino acid chain is forest green and the glycosylations are magenta.

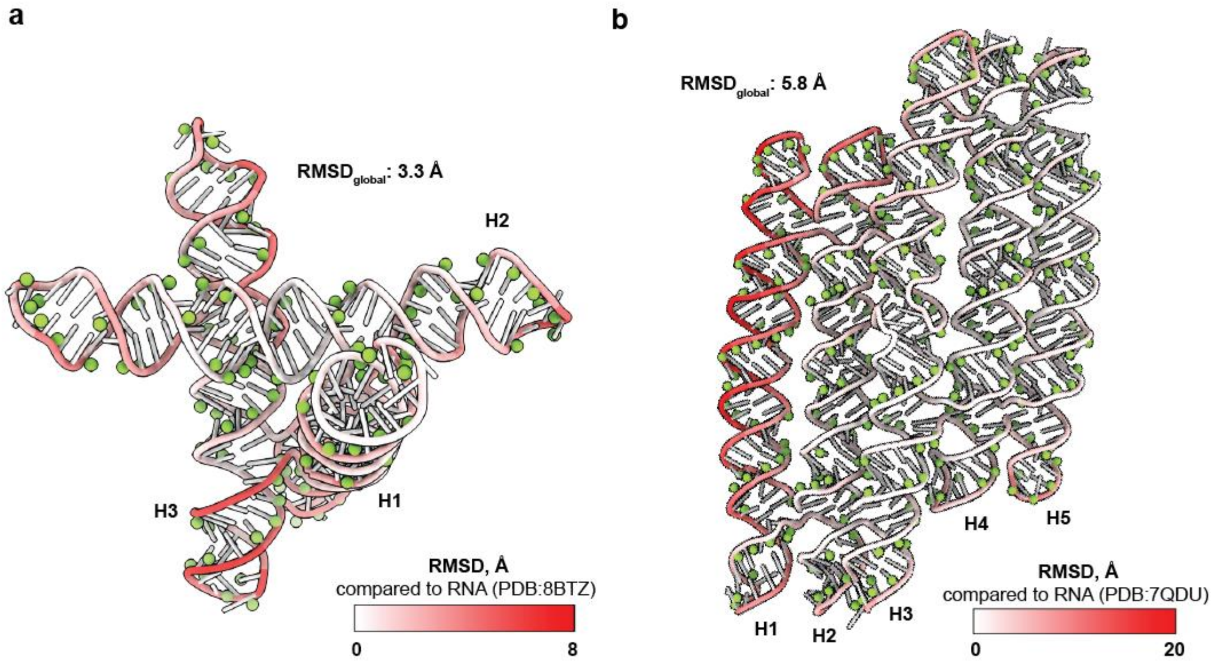

**Figure E4. Comparison between cryo-EM derived models for RNA and FY-RNA origamis.** The cryo-EM informed models of the FY-RNA PXT (**a**) and 5HB (**b**) structures are compared with their already published RNA versions by RMSD (see scale bar). F-atoms are shown as green spheres.

>6HBC-dimer

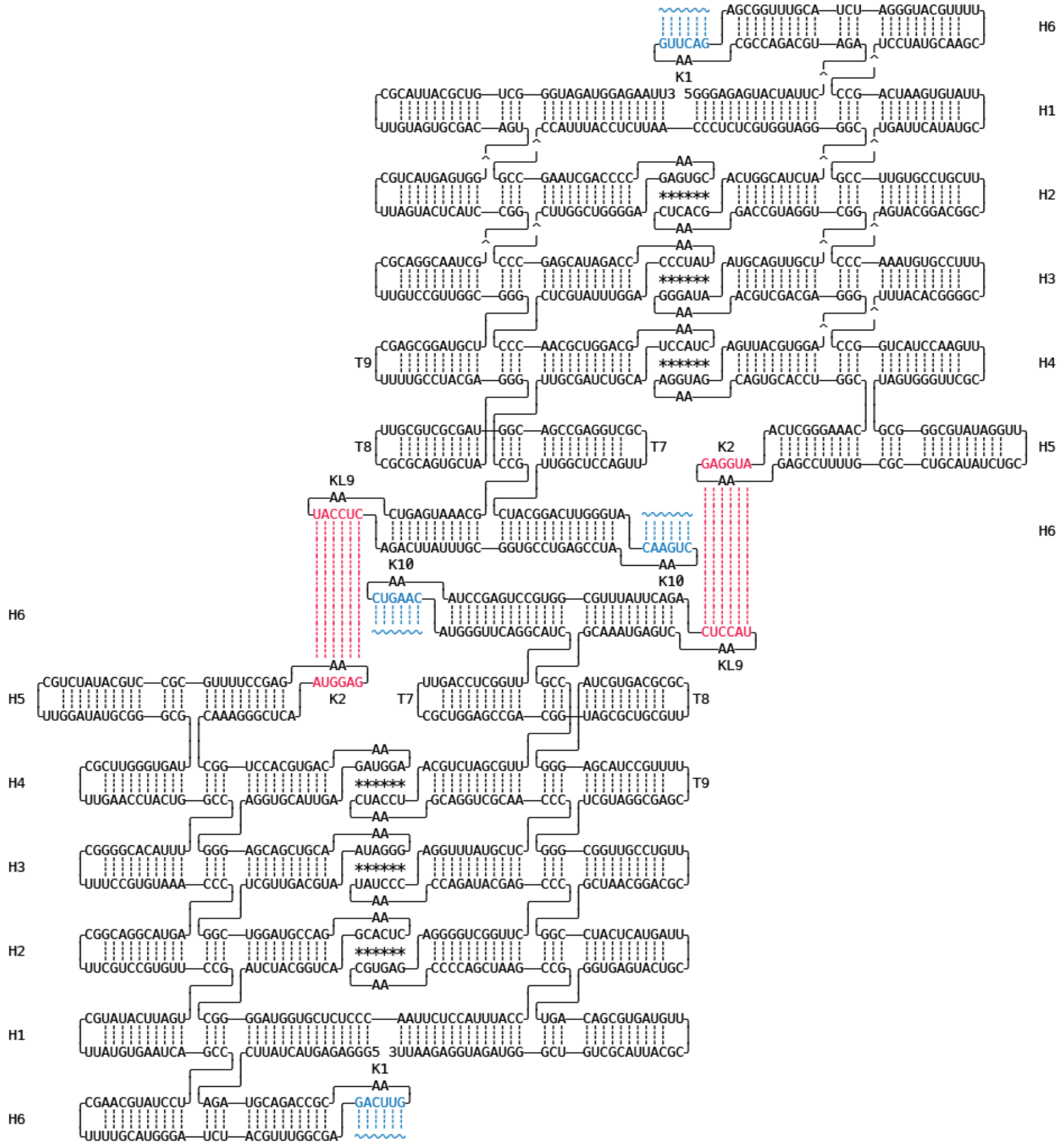

**Figure E5. Quaternary structure of 6-helix bundle clasp dimer (6HB-dimer).** Coaxially stacked helices (H) are numbered from 5' to 3' end except for H6. Kissing loop (KL) base pairs are annotated by *. KLs and TLs are numbered from 5' to 3' end. Intermolecular base pairs between KL2 and KL9 are marked in red. Intramolecular base pairs between KL1 and KL10 are marked in cyan interrupted by a ~ symbol. KL1, KL2, KL9 and KL10 are annotated next to their AA-bulges. TL7-9 are annotated.

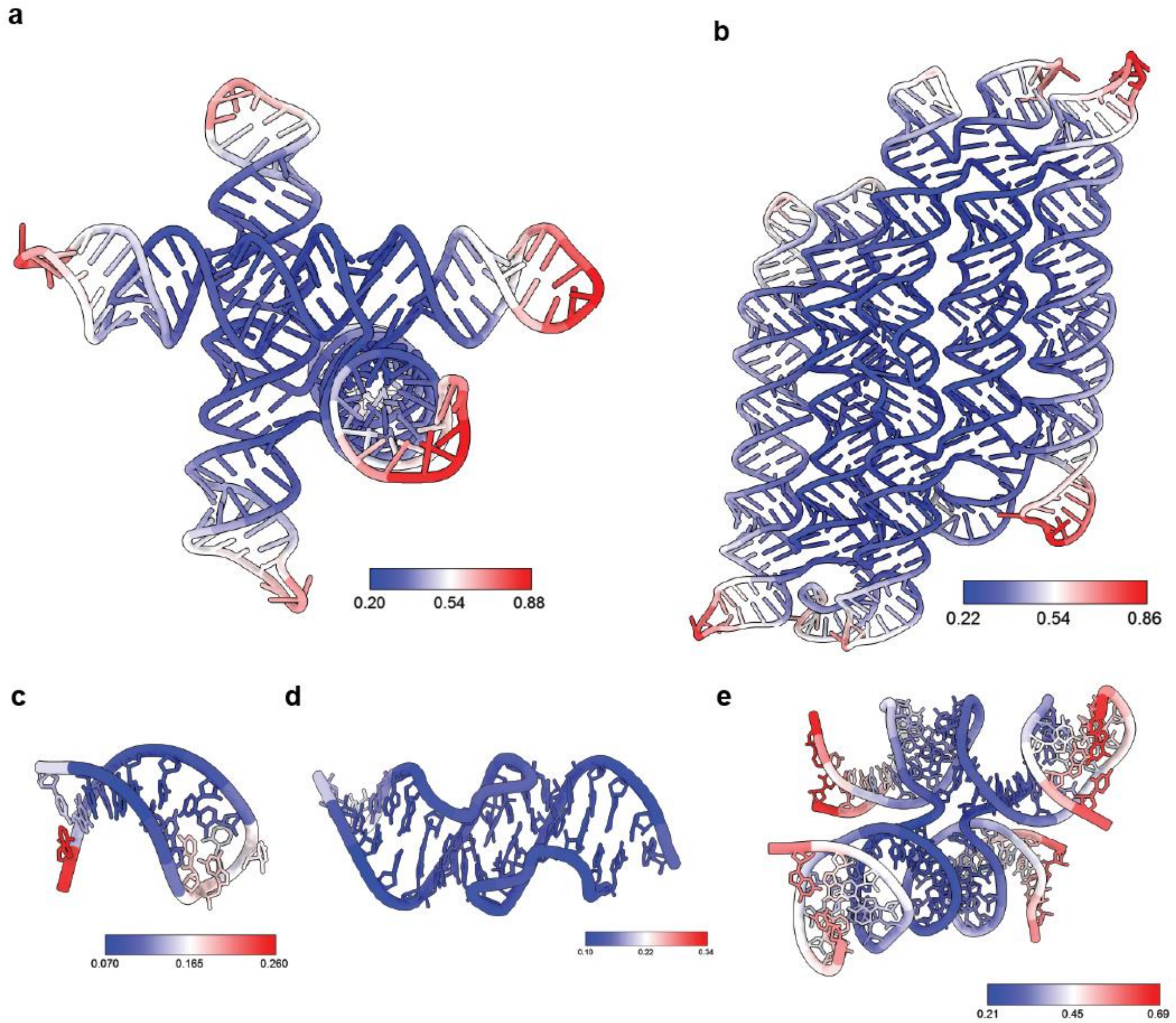

**Figure E6. RMSF of simulated FY-RNA origamis. a-e**, MD simulated models described in the main text coloured locally in relation to their local root mean square fluctuation (RMSF). Scale bar values denote RMSF in nanometres (nm).

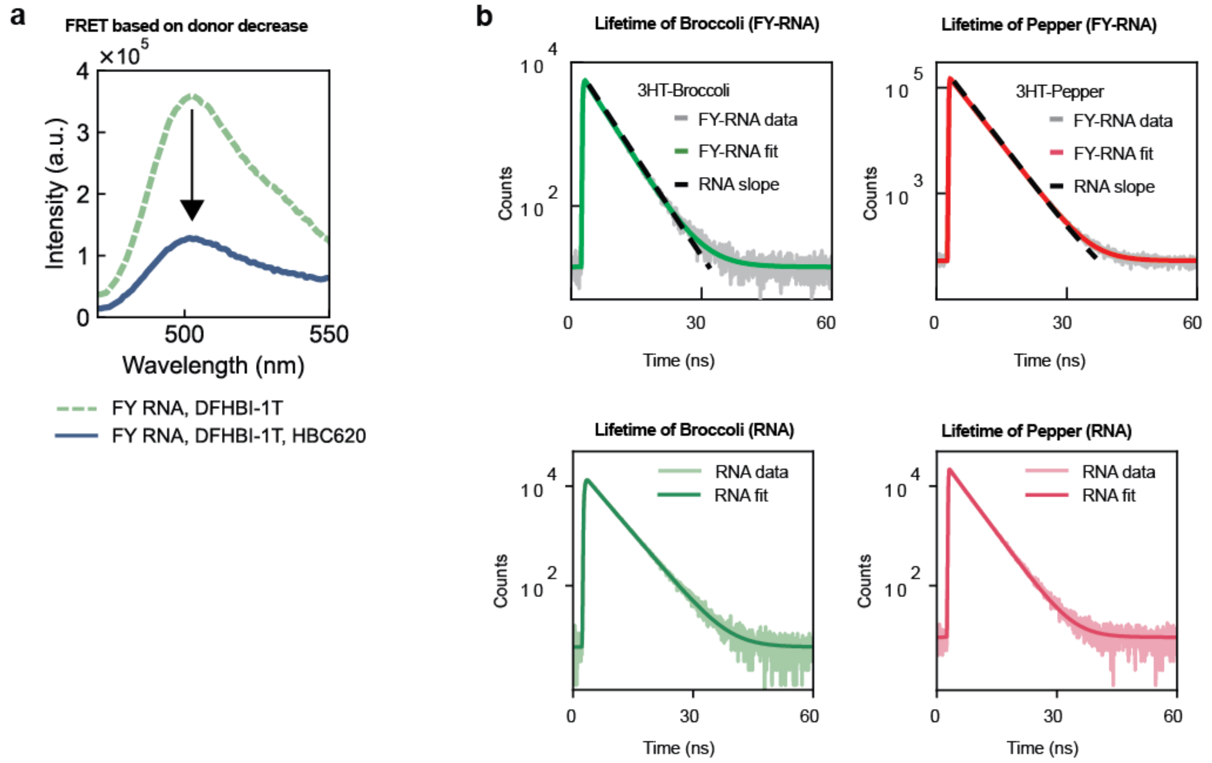

**Figure E7. FRET and fluorescence lifetime measurements. a**, Fluorescence spectrum of FY-RNA 3HT-BP with DFHBI-1T (green, dashed curve) or DFHBI-1T and HBC620 (blue, solid curve), showing a decrease in DFHBI-1T emission in presence of HBC620, indicating FRET. **b**, Fluorescence lifetime measurement of the 3HT-BP as RNA and FY-RNA. Broccoli was measured by adding DFHBI-1T and Pepper was measured by adding HBC to the samples. Fluorescence lifetime measurement of 3HT-Broccoli or 3HT-Pepper showing that the decay of FY-RNA (coloured lines) is similar that of RNA (dashed lines), indicating that the environment for fluorescence is similar in the subfraction of particles that fluoresce.

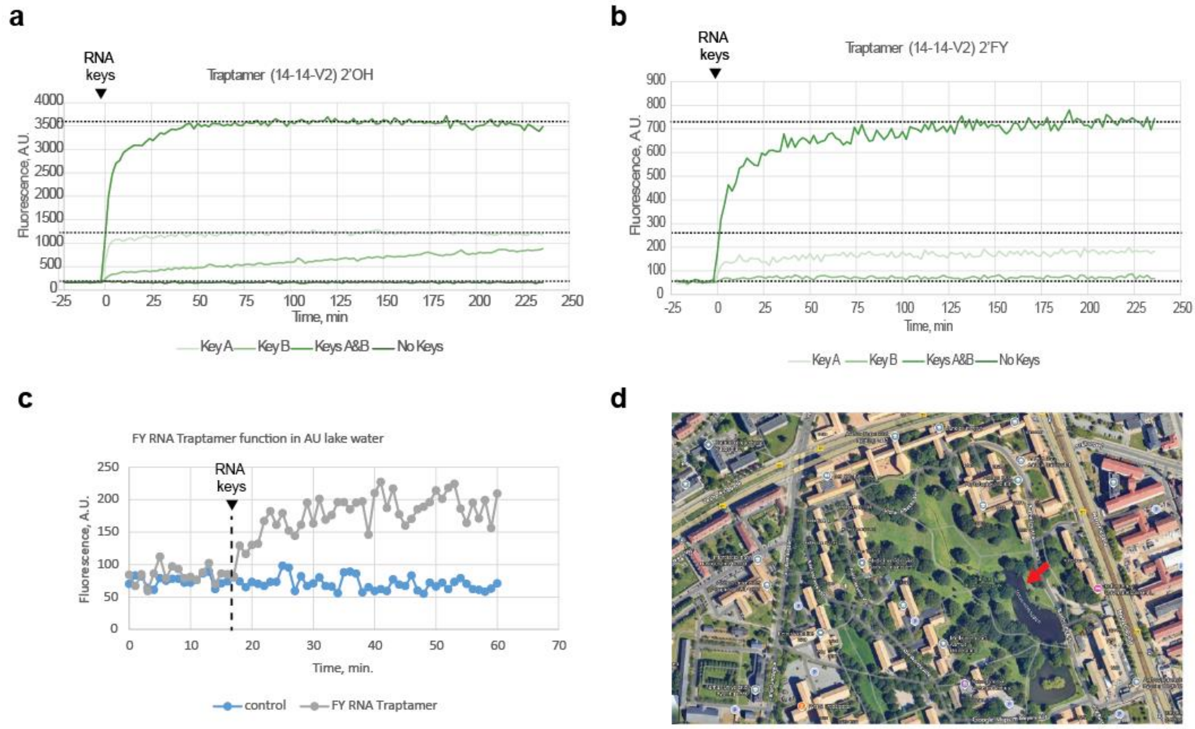

**Figure E8. Traptamer AND gate function and stability lake water. a-b**, Fluorescence experiments to test AND gate function for RNA and FY-RNA Traptamer device. **c**, Function of FY-RNA Traptamer in Aarhus University Lake water. **d**, Arrow point to the lake in the Aarhus University campus park. Lake samples were retrieved after the renowned boat race event "Kapsejladsen".

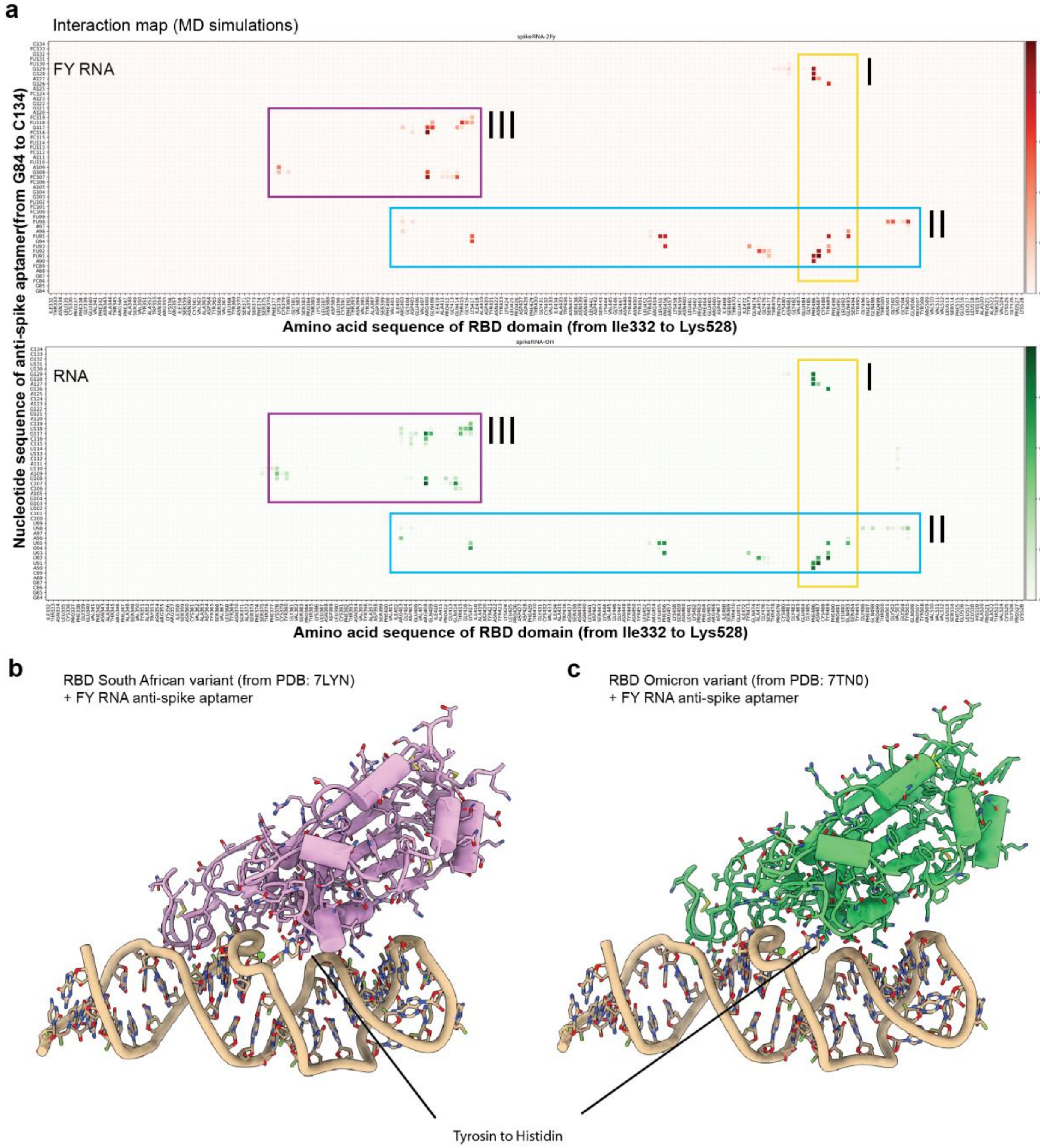

**Figure E9. a**, MD simulation interactions plotted as amino acids on the x-axis and nucleotides on the y-axis. **b-c**, Model of SA and Omicron RBDs positioned together with the anti-Spike aptamer. RBD were aligned using ChimeraX.

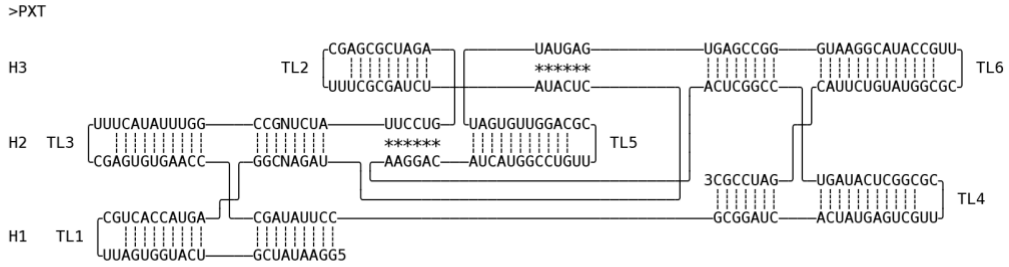

**Figure S1. Blueprint for paranemic crossover tile (PXT).** Coaxially stacked helices (H) are numbered from 5' to 3' end. Paranemic crossover (PX) base pairs are annotated by *. TLs are numbered from 5' to 3' end. TL3 is marked.

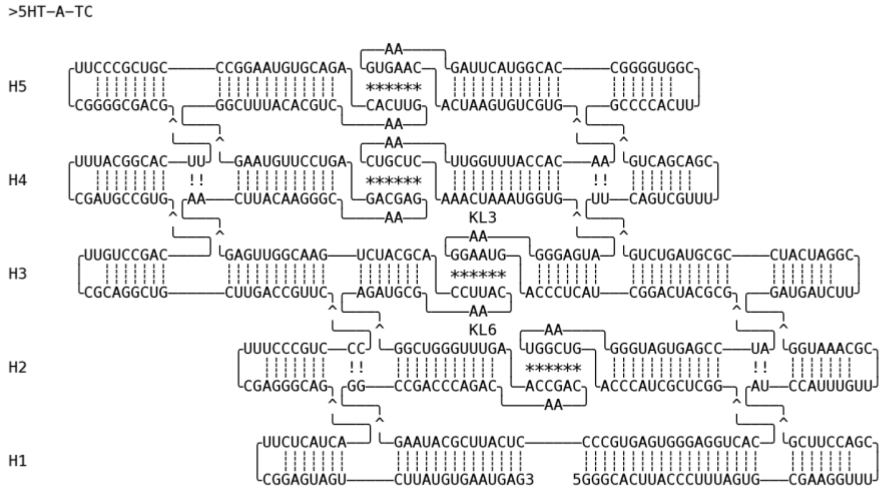

**Figure S2. Blueprint for 5-helix tile A twist corrected (5HT-A-TC).** Coaxially stacked helices (H) are numbered from 5' to 3' end. Kissing loop (KL) base pairs are annotated by *. KL3 and KL6 are annotated next to their AA-bulges. Dovetail base pairs are annotated with !.

>6HBC

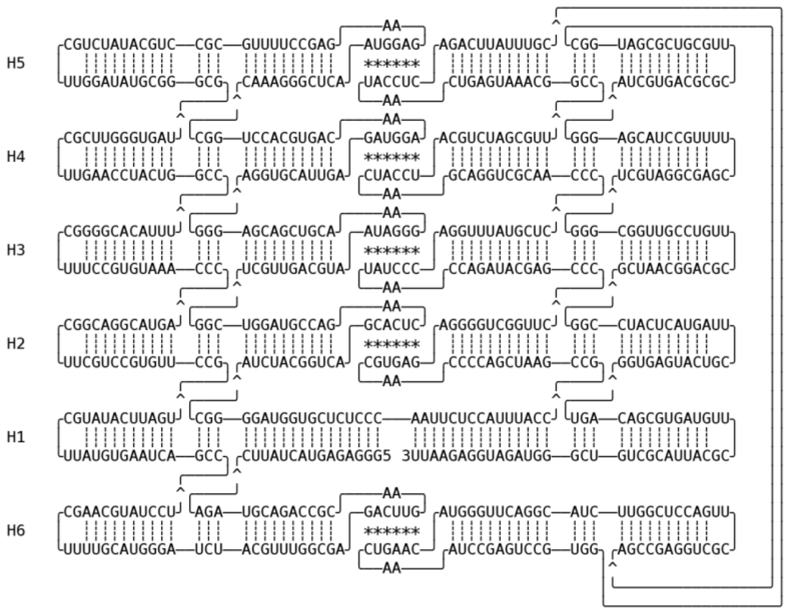

**Figure S3. Blueprint for 6-helix bundle clasp (6HBC).** Coaxially stacked helices (H) are numbered from 5' to 3' end except for H6. Kissing loop (KL) base pairs are annotated by *.

>6HBC-TC

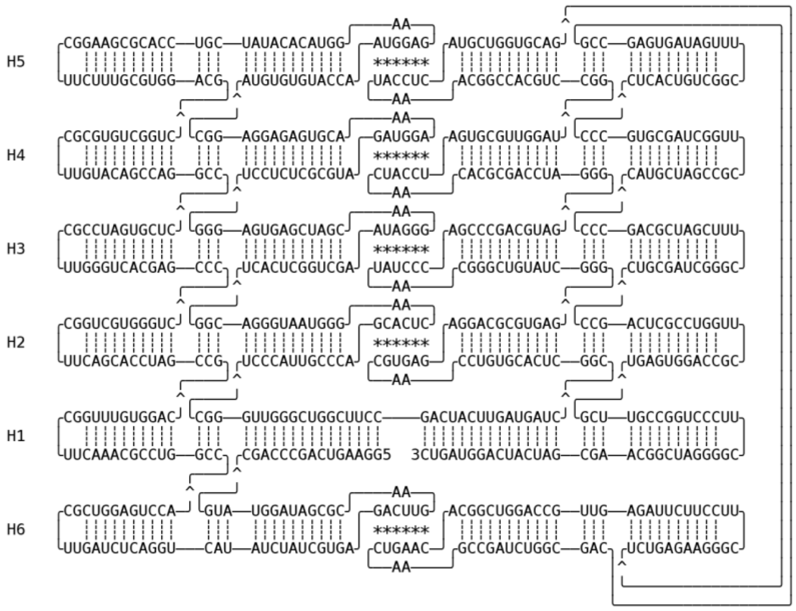

**Figure S4. Blueprint for 6-helix bundle clasp twist corrected (6HBC-TC).** Coaxially stacked helices (H) are numbered from 5' to 3' end except for H6. Kissing loop (KL) base pairs are annotated by *.

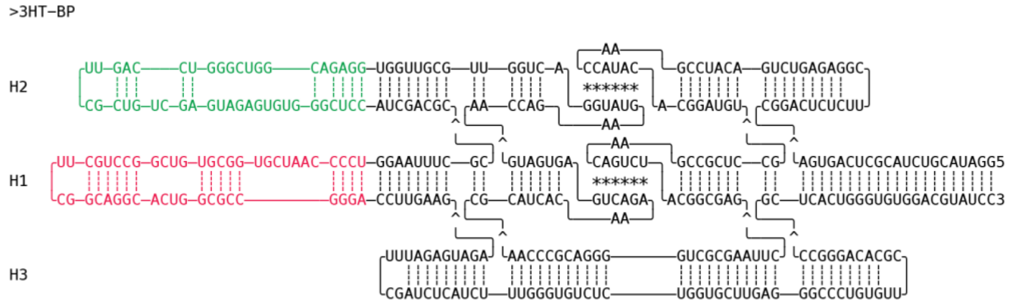

**Figure S5. Blueprint for 3-helix tile with Broccoli and Pepper aptamers (3HT-BP).** Coaxially stacked helices (H) are numbered from 5' to 3' end. Kissing loop (KL) base pairs are annotated by *. Broccoli marked in green, Pepper in red.

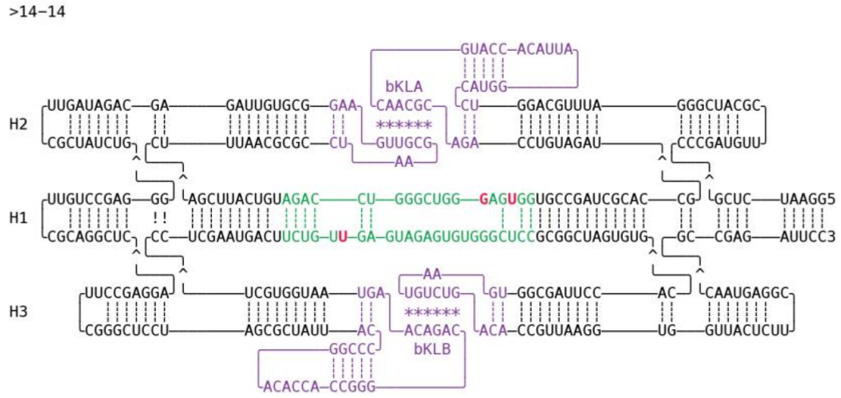

**Figure S6. Blueprint for 3-helix tile with internal iSpinach aptamer (Traptamer).** Coaxially stacked helices (H) are numbered from 5' to 3' end. Branched kissing loop (bKL) base pairs are annotated by * and annotated as bKLA and bKLB (purple). The sequence differences in the iSpinach aptamer (green) in comparison to the Broccoli aptamer are marked in bold and red.

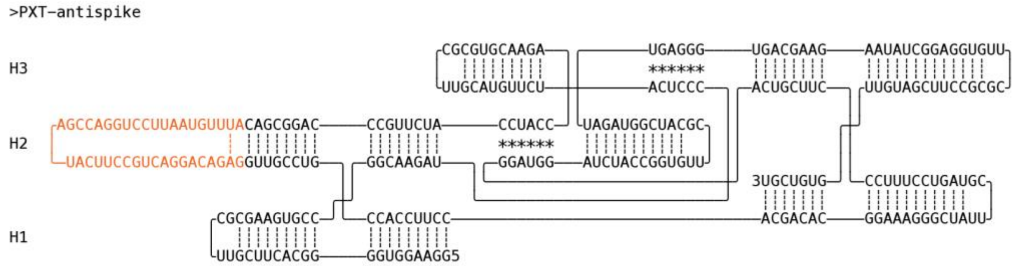

**Figure S7. Blueprint for paraneic crossover tile with anti-Spike aptamer (PXT-Spike).** Coaxially stacked helices (H) are numbered from 5' to 3' end. Paranemic crossover (PX) base pairs are annotated by *. The anti-Spike aptamer is marked in orange.

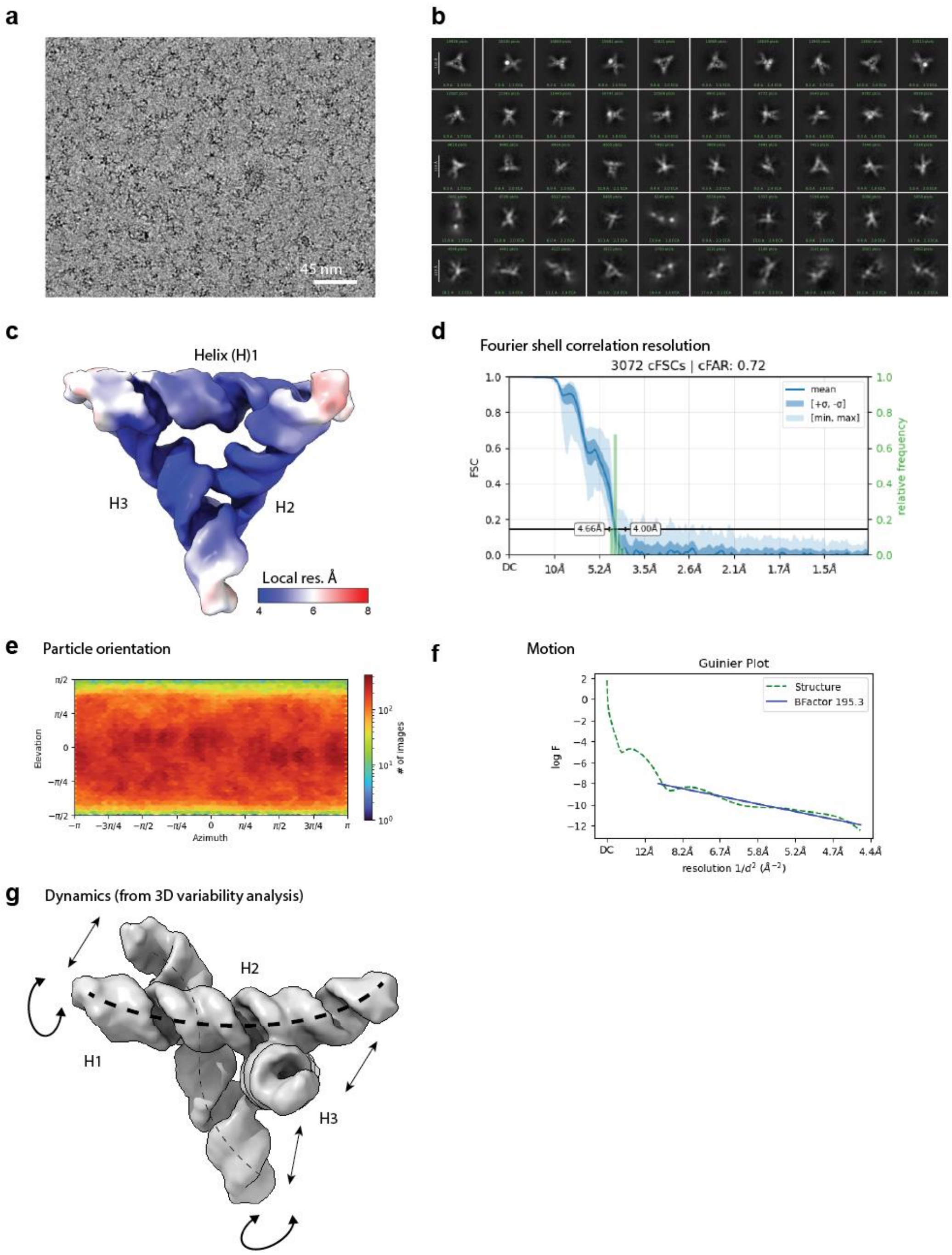

**Figure S8. Cryo-EM analysis overview of the FY-RNA paranemic crossover triangle (PXT).** Examples of (**a**) contrast transfer function (CTF)-fitted and motion corrected micrograph of ice with FY-RNA PXT particles and (**b**) 2D classes generated during the cryo-EM analysis. **c**, show cryo-EM density coloured for local resolution, (**d**) the Fourier shell correlation (FSC) plot, (**e**) the particle orientation, and (**f**) the B-factor fit. **g**, shows the observed density variation assessed by 3D classification and 3D variability analysis indicative of internal particle dynamics and flexibility. Arrows and dashed lines indicate the twisting and bending of the helices relative to each other.

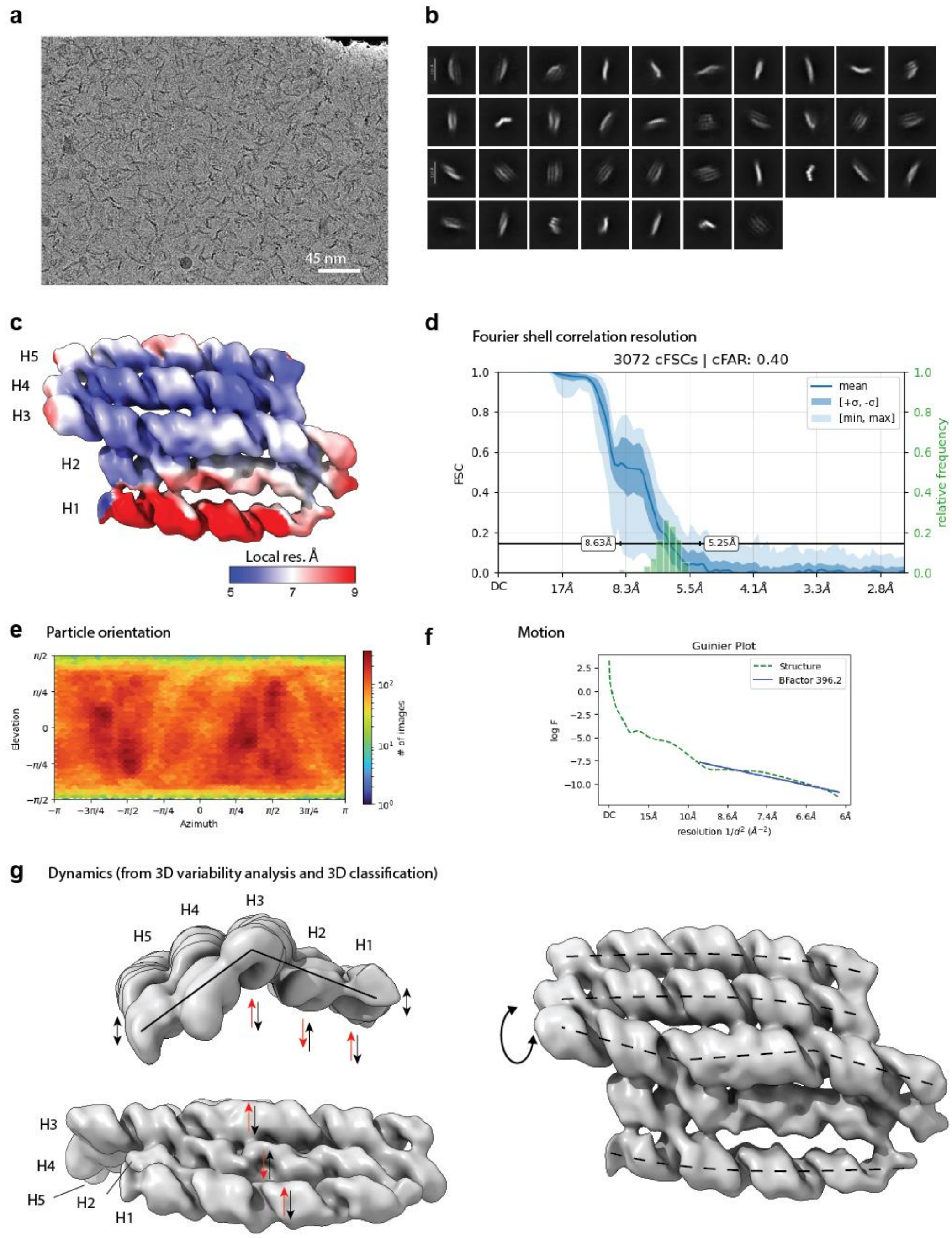

**Figure S9. Cryo-EM analysis overview of the FY-RNA 5-helix tile (5HT).** Examples of (**a**) contrast transfer function (CTF)-fitted and motion corrected micrograph of ice with FY-RNA 5HT particles and (**b**) 2D classes generated during the cryo-EM analysis. **c**, show cryo-EM density coloured for local resolution, (**d**) the Fourier shell correlation (FSC) plot, (**e**) the particle orientation, and (**f**) the B-factor fit. (**g**) shows the observed density variation assessed by 3D classification and 3D variability analysis indicative of internal particle dynamics and flexibility. Arrows and dashed lines indicate the twisting and bending of the helices relative to each other. Red and black arrows indicate that helices bend up and down in turn.

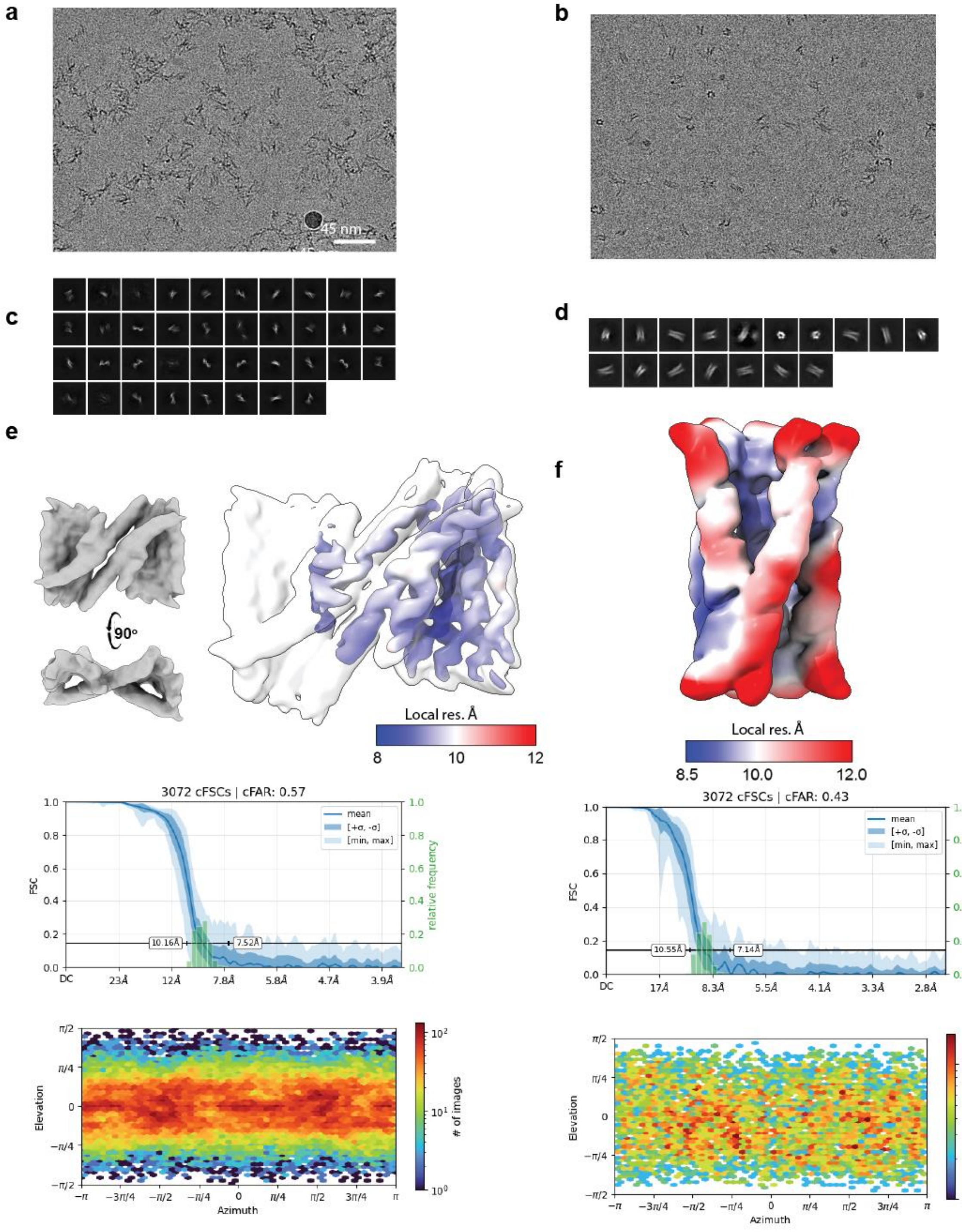

**Figure S10. Cryo-EM analysis overview of the FY-RNA 6-helix bundle with a clasp (6HB-C). a,b**, Examples of contrast transfer function (CTF)-fitted and motion corrected micrograph of ice with two different FY-RNA 6HB-C designs, the optimized design and the original design, respectively. **c,d**, show 2D classes generated during the cryo-EM analysis of the respective samples. **e,f**, show cryo-EM density coloured for local resolution, Fourier shell correlation (FSC) plots, and particle orientation plots.

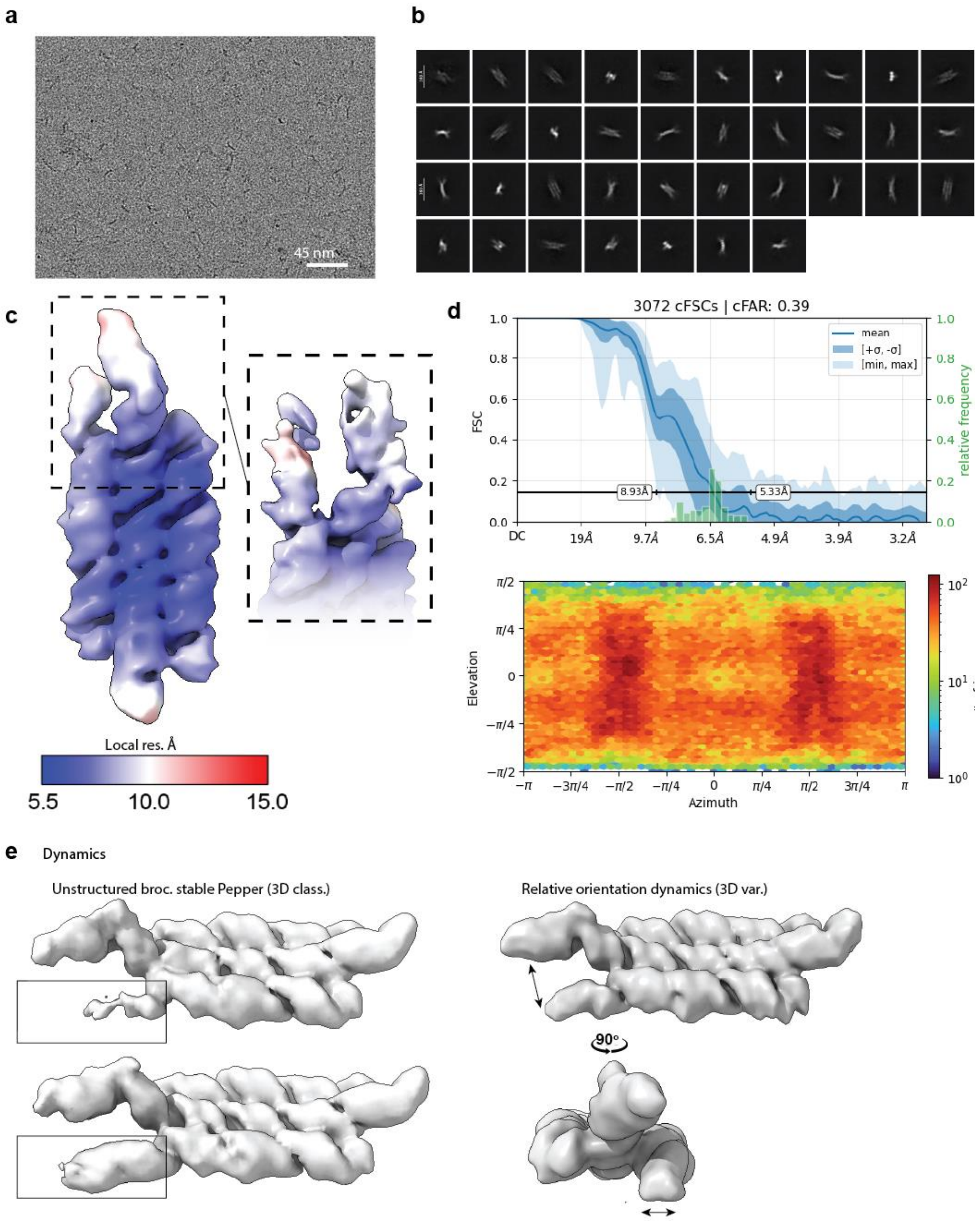

**Figure S11. Cryo-EM analysis overview of the FY-RNA 3-helix tile with Papper and Broccoli (3HT-PB). a**, Example of contrast transfer function (CTF)-fitted and motion corrected micrograph of ice with FY-RNA 3HT-PB particles. **b**, 2D classes generated during the cryo-EM analysis. **c**, Cryo-EM density coloured for local resolution. **d**, Fourier shell correlation (FSC) plot and orientation analysis. **e**, Result of 3D classification analysis (left) revealing sub-populations of the particles where Broccoli is undefined in comparison with the rest of the structure. Results of the 3D variability analysis (right) revealing internal dynamics in the relative positioning and orientation of the fluorogenic aptamers.

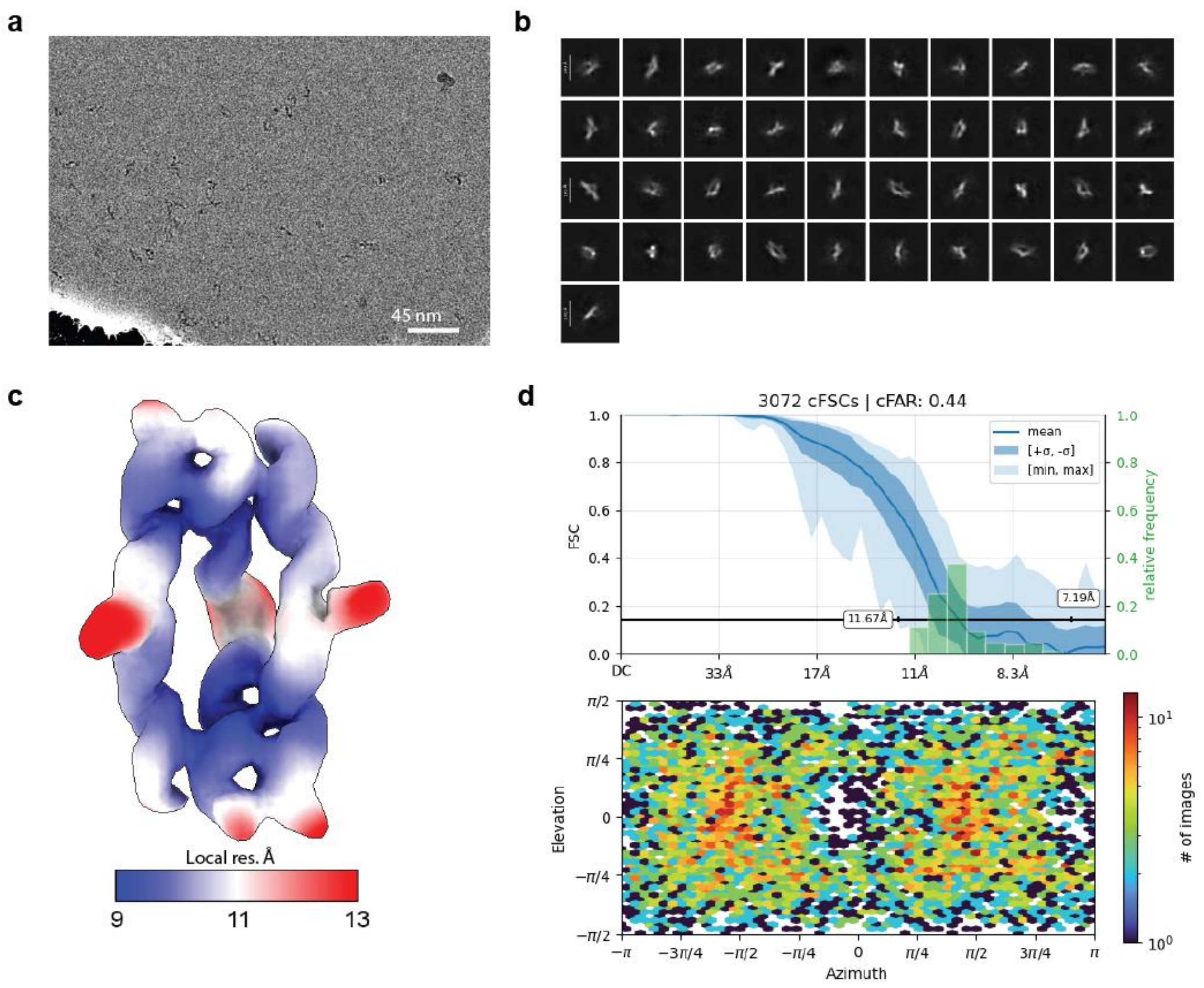

**Figure S12. Cryo-EM analysis overview of the FY-RNA Traptamer. a**, Example of contrast transfer function (CTF)-fitted and motion corrected micrograph of ice with FY-RNA Traptamer particles. **b**, 2D classes generated during the cryo-EM analysis. **c**, Cryo-EM density coloured for local resolution. **d**, Fourier shell correlation (FSC) plot and orientation analysis.

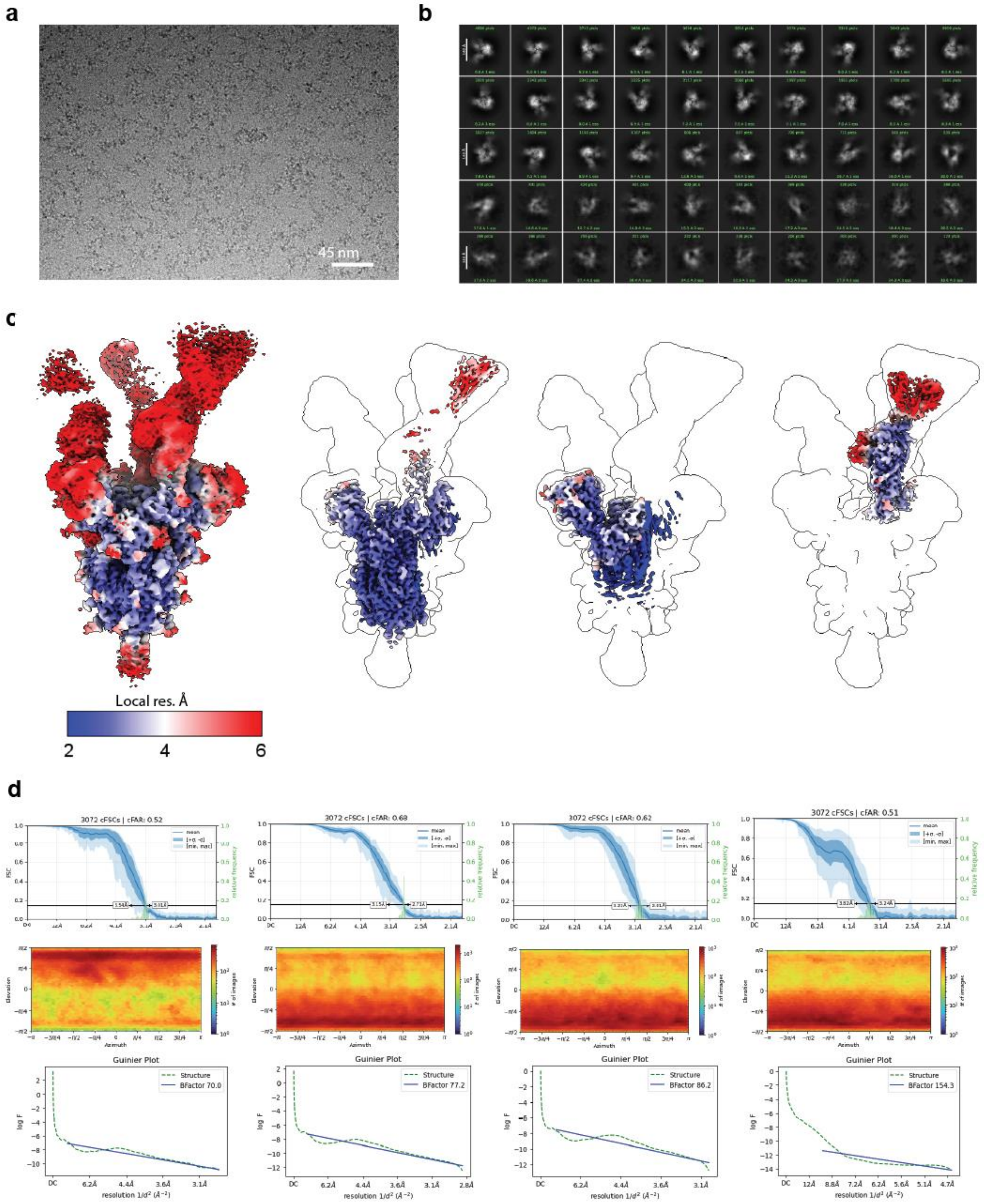

**Figure S13. Cryo-EM analysis overview of the SARS-CoV-2 spike protein with FY-RNA PXT with anti-spike protein aptamer. a**, Contrast transfer function (CTF)-fitted and motion corrected micrograph of ice with SARS-CoV-2 spike protein and FY-RNA PXT with anti-spike protein aptamer particles. **b**, 2D classes generated during the cryo-EM analysis. **c**, Cryo-EM density coloured for local resolution of the different focus maps. **d**, Fourier shell correlation (FSC) plot, orientation analysis and Guinier plots.

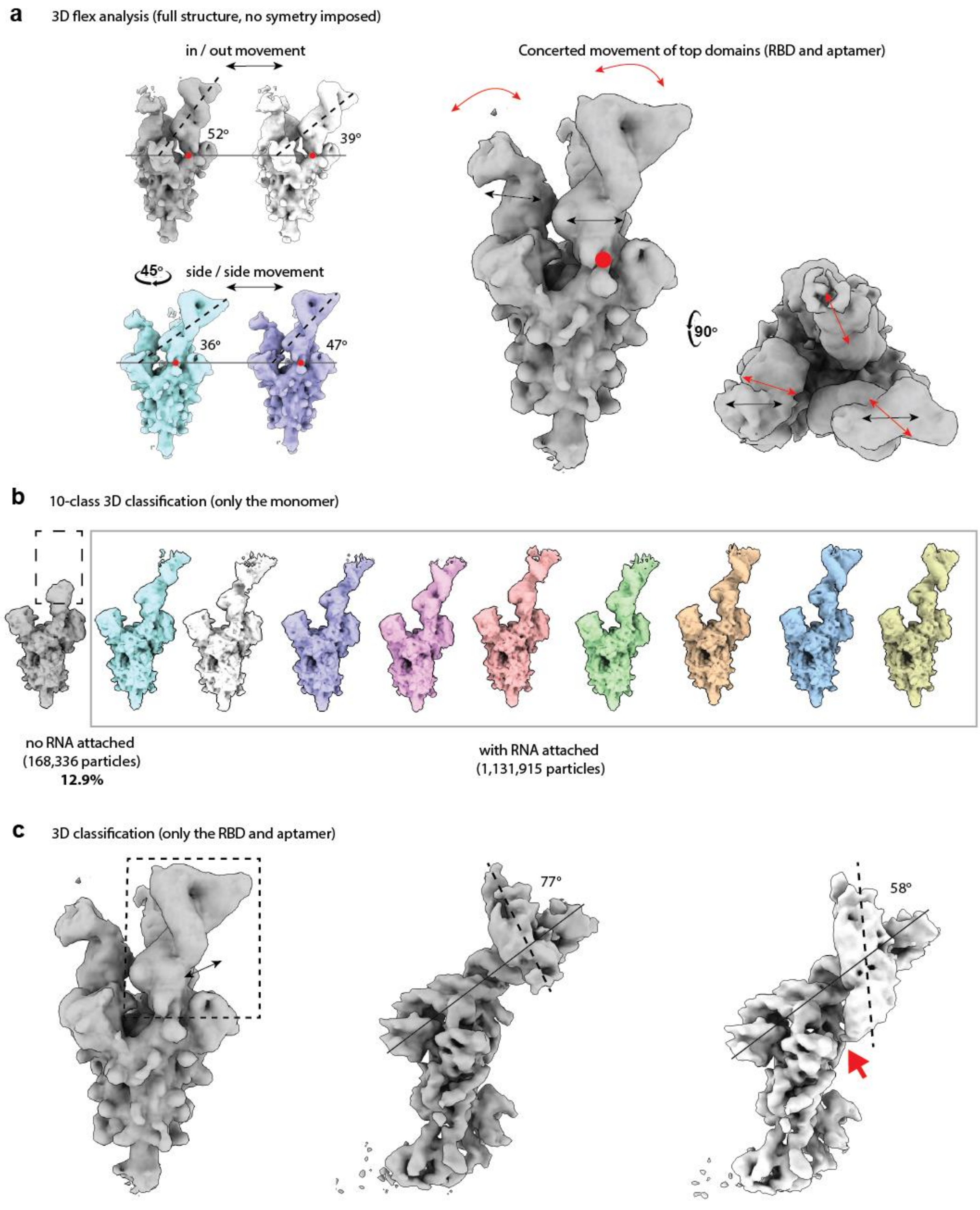

**Figure S14. Dynamic analysis of the SARS-CoV-2 spike protein with FY-RNA PXT with aptamer density maps. a**, 3Dflex analysis of the non-symmetry expanded structure revealing coordinated movement of the regions corresponding to the RBD and FY-RNA origami devise is located. The movement happens around a "hinge point" (red spot) at the base of the RBD domain. This concerted movement and the extra density at all three RBD locations, indicate that at least some of the particles have three PXT with aptamers bound simultaneously. However, the extra density at the RBD location is strongest at position 1 and diminishes at positions 2 and 3, indicating that not all particles have three FY-RNA origami devices bound, some have two and some have one. **b**, 10-class 3D classification of symmetry expanded particles revealing that 12.9 % did not have extra density at the RBD domain. This likely represents the fraction of particles that do not have any FY-RNA origami device bound. Assuming no cooperativity, this would mean that 66.1% of the spike proteins have three origamis bound, 29.4% would have two, and 4.3% would have one, and 0.2% would have no bound. **c**, 3D classification of the RBS focus map, revealing the same kind of dynamic movement observed in (A), and in addition some dynamics of the angle of the PXT helix 1, sometimes interacting with the RBD-domain through the TL (red arrow).

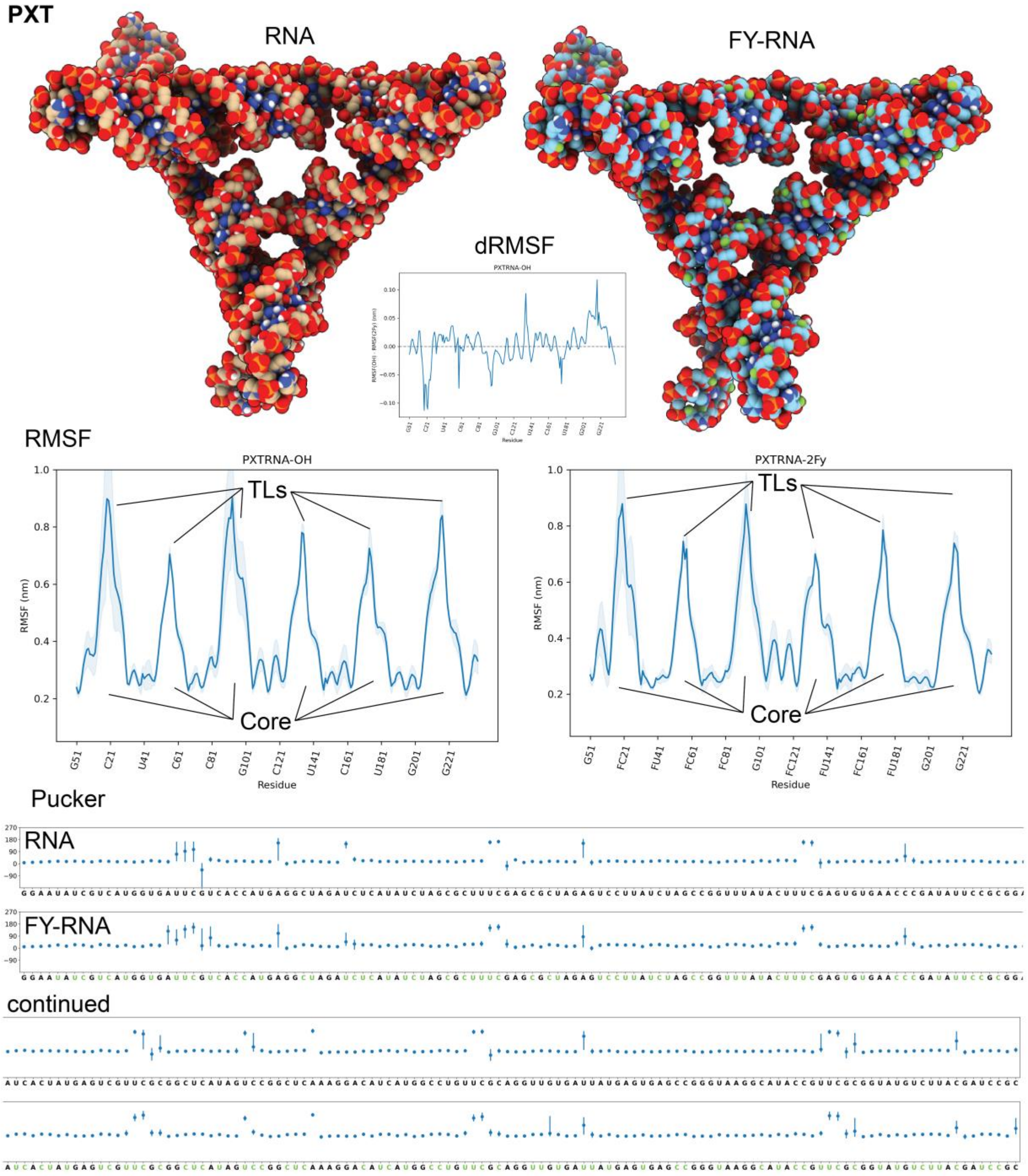

**Figure S15. MD-simulation of the PXT.** Simulated structures of the PXT as RNA and FY-RNA. Corresponding RMSF, ΔRMSF, and Pucker plots are shown. The simulation reveals that fluctuation was observed primarily in the six helix ends, indicated by the six peaks in RMSF. The pucker plot shows that C2'-endo conformation was seen in the U and C residues of the UUCG TL and sometimes in the 5'-end residues of the crossover (indicated by arrows).

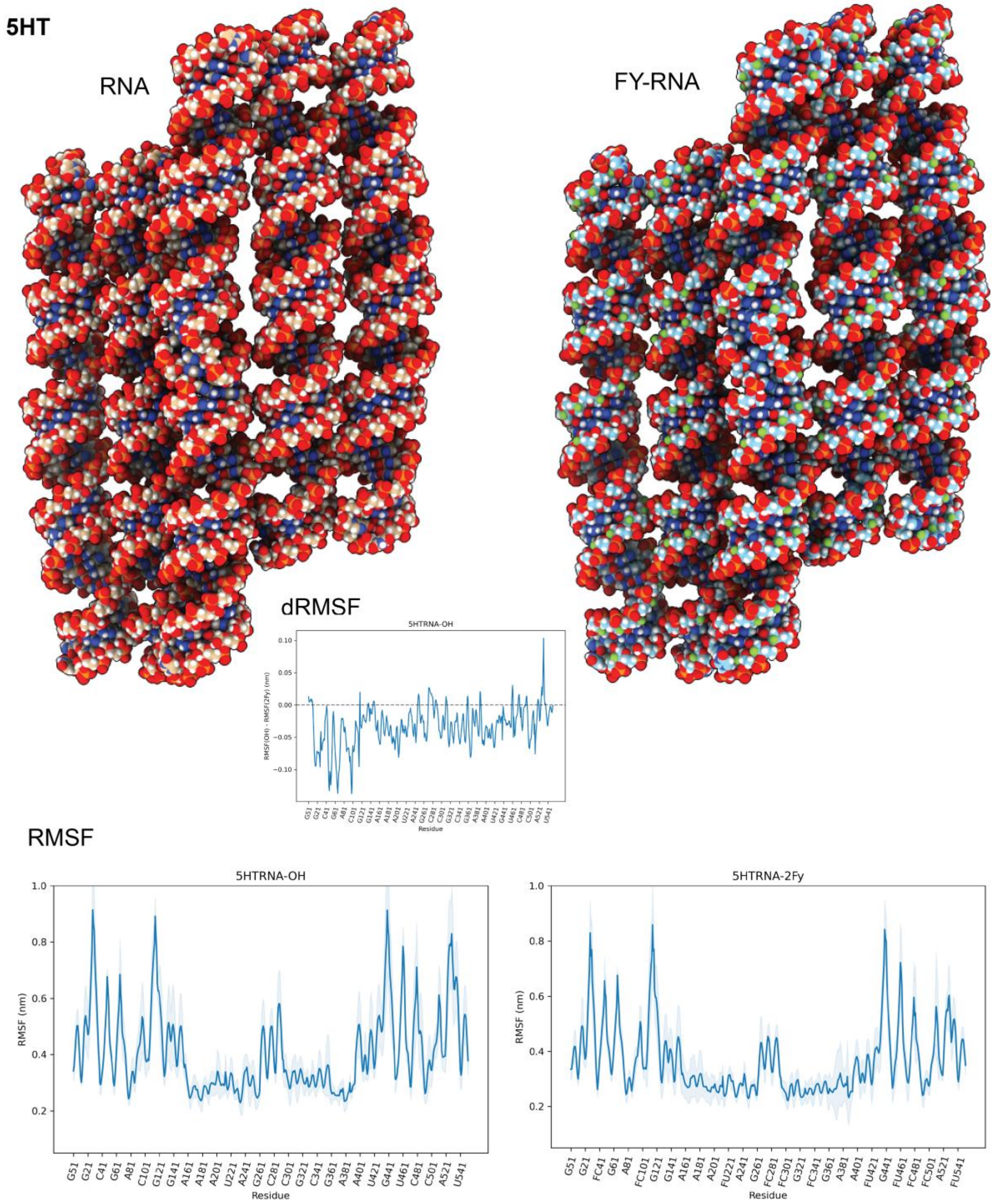

**Figure S16. MD-simulation of the 5HT.** Simulated structures of the 5HT as RNA and FY-RNA. Corresponding RMSF and ΔRMSF plots are shown. The simulation reveals that fluctuation was observed in the 10 helix ends, indicated by the peaks in RMDF in the start and end of the sequence. In addition, the central part of the molecule (between ~G261 and C301) was fluctuating. This is the region that base pairs with the 5'- and 3'-end of the molecule.

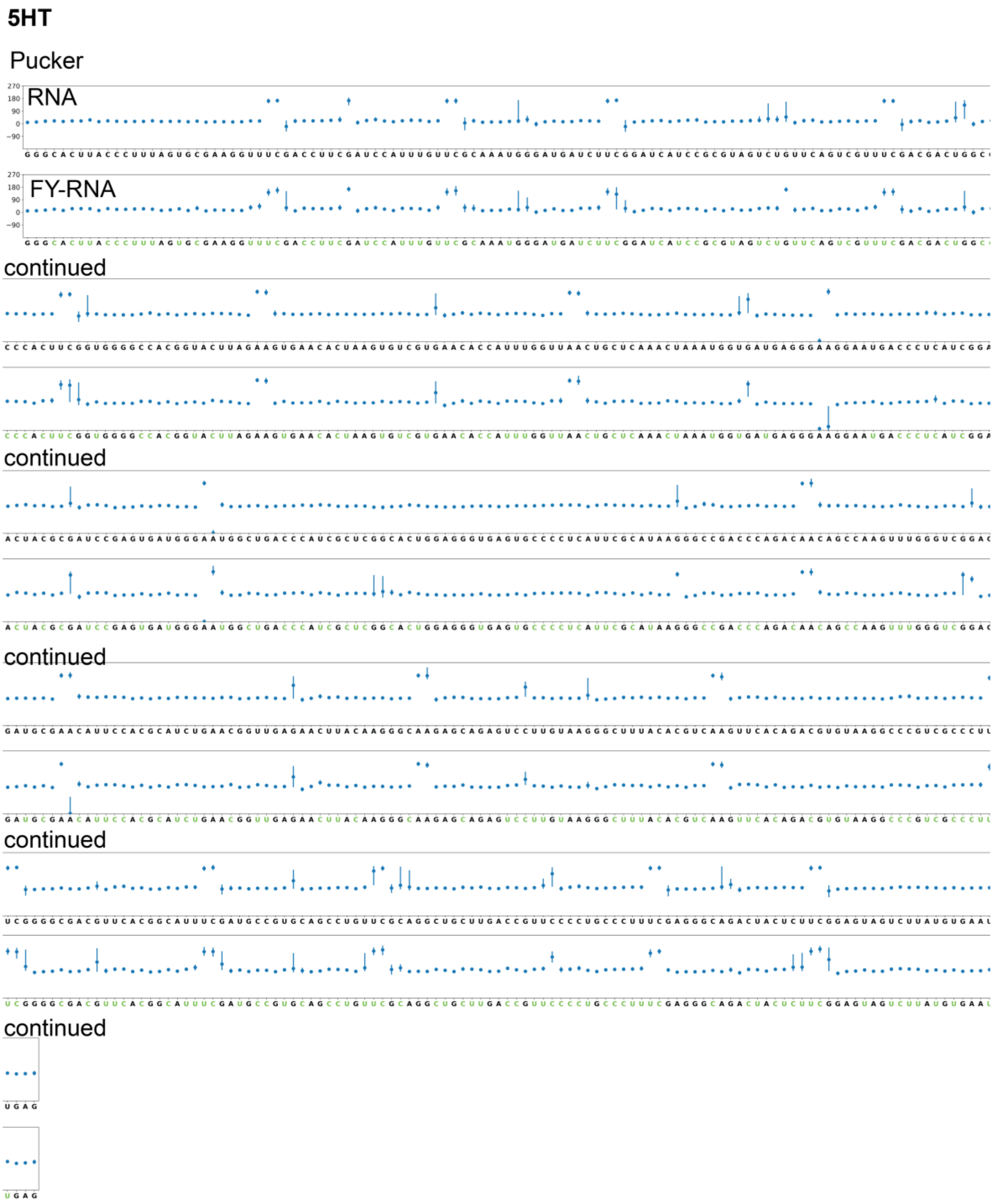

**Figure S17. Pucker plots for the 5HT.** Simulated structures of the 5HT as RNA and FY-RNA. The pucker plot shows that C2'-endo conformation was seen in the U and C residues of the UUCG TL, the A-residues of the KLs, and sometimes in the 5'-end residues of the crossover (indicated by arrows).

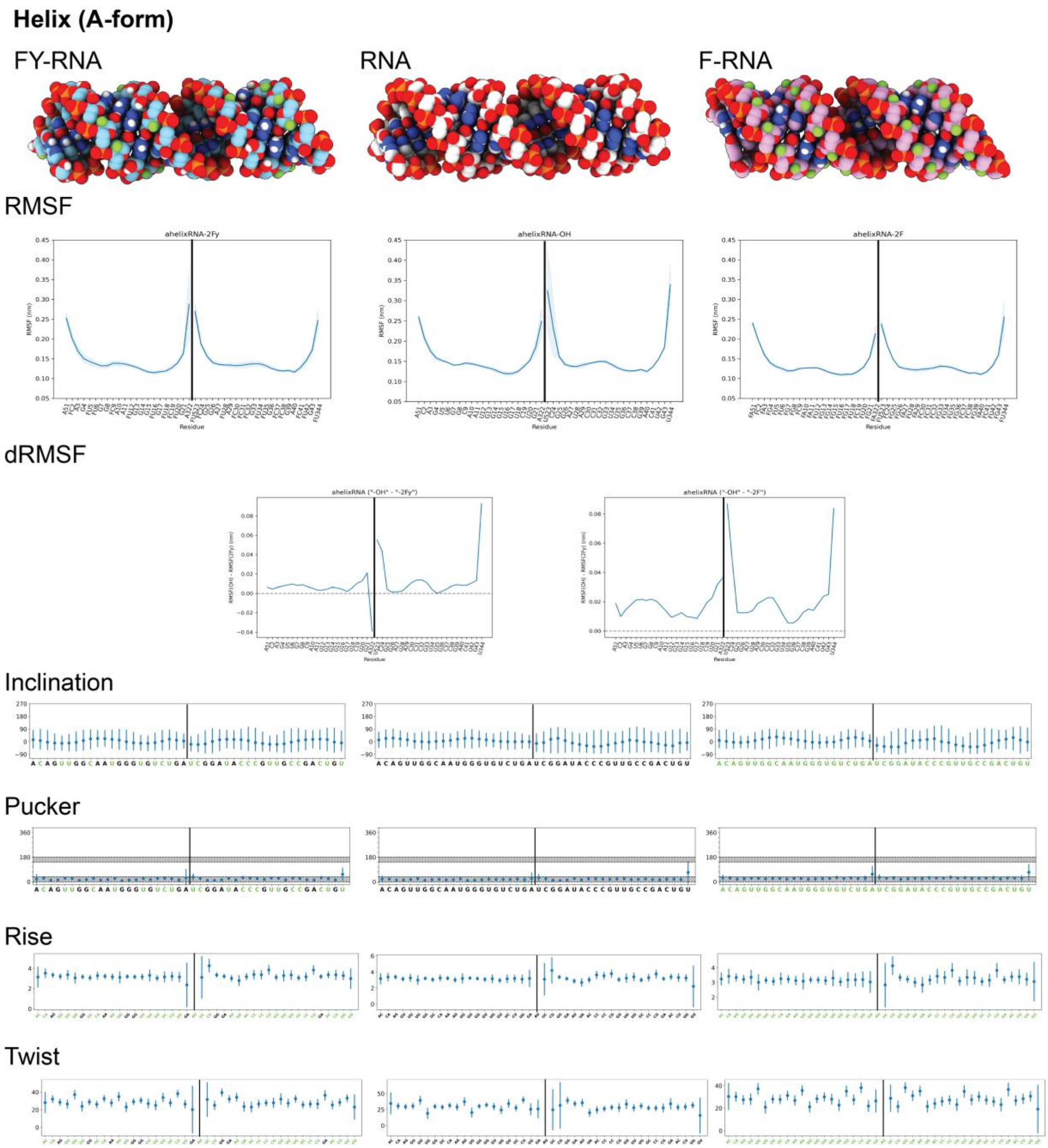

**Figure S18. MD-simulation of A-form helical NAs.** Simulated structures of double helix segment of FY-RNA, RNA and F-RNA and corresponding RMSF, ΔRMSF, Inclination, Pucker, Rise and Twist plots. These look very similar as expected for the A-from helix and is used as a validation of the forcefield and parameters of the simulation.

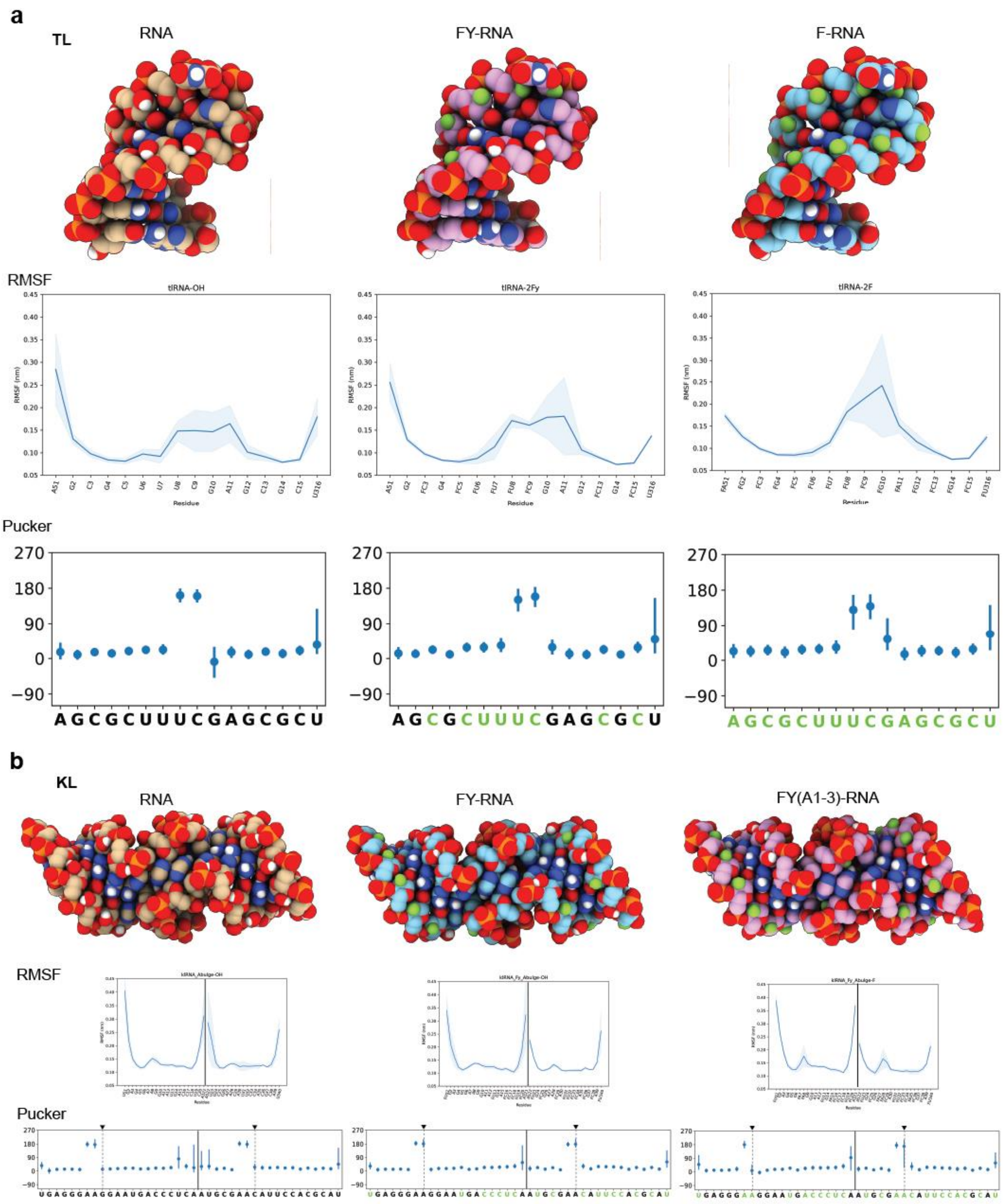

**Figure S19. MD-simulation of the TL (top) and KL (bottom). a**, Simulated structures of the UUCG TL as RNA, FY-RNA and F-RNA. Corresponding RMSF, and Pucker plots are shown. The simulation reveals that fluctuation increases primarily in the G residue of the UUCG TL when it is simulated as FY-RNA or F-RNA. The pucker plots show that C2'-endo conformation is prominent in the U and C residues of the UUCG TL (indicated by arrows). This fits with the observation from the simulations of the full PXT and 5HT origamis. **b**, Simulated structures of the A-bulge KL as RNA, FY-RNA and FY-RNA + FY(A1-3)-RNA. Corresponding RMSF, and Pucker plots are shown. The simulation reveals a peak in the fluctuation of a particular A-bugle As only if these are simulated as F-RNA. The pucker plots show that the A-bulge A are almost exclusively in C2'-endo conformation, but that when changed to F-RNA this is changed. This fits with the observation from the simulations of the whole origamis PXT and 5HT showing that FY-RNA KLs are folding fine, but that F-RNA As are destabilized. Simulation of the FY-RNA + F-RNA A-bulge As KL is, however, still stable and might form.

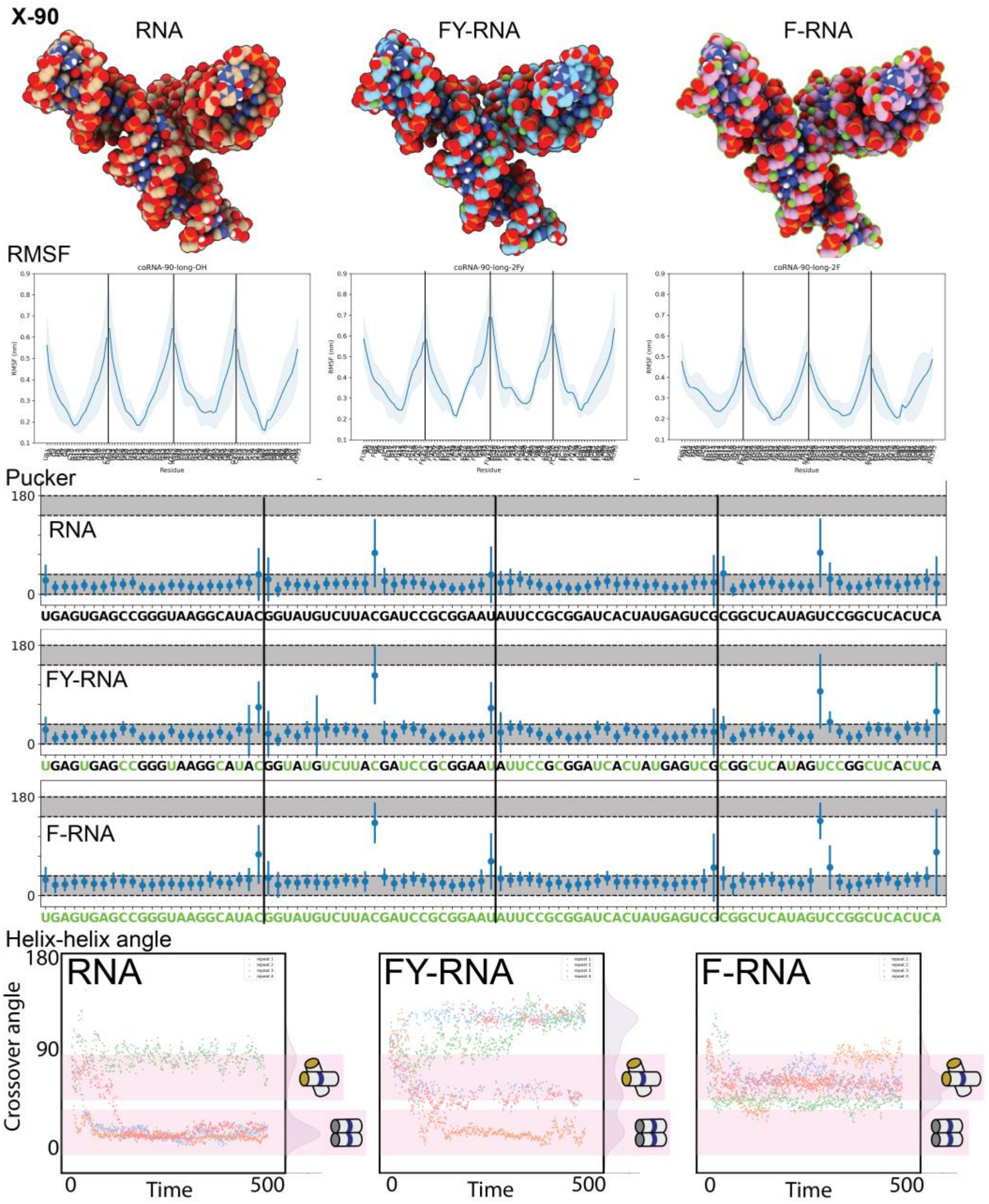

**Figure S20. MD-simulation of the X-90 crossover.** Simulated structures of the X-90 as RNA, FY-RNA and F-RNA. Corresponding RMSF, Pucker plots, and helix angle plots are shown. The simulation reveals that fluctuation is lowest at the centre of the strands, which is where the CO is positioned. The pucker plots show that C2'-endo conformation is sometimes adapted in the 5' residue of the CO (indicated by arrows). Surprisingly the most adaptation of C2'-endo conformation is seen for the F-RNA, and could be explained by the increased base paring in the helixes (compared to RNA) which forces the less favourable conformation in the backbone of the residue that facilitates the CO. This fits with the observation from the simulations of the whole origamis PXT and 5HT. The Helix angle plot shows that the simulation of the RNA X-90 in most cases makes a parallel interaction, whereas FY-RNA and F-RNA adopts a crossed conformation, and remain to be more dynamic throughout the simulations. The parallelization of the RNA X-90 is caused by formation of HH interactions which are perturbed when the 2'-position is fluorinated.

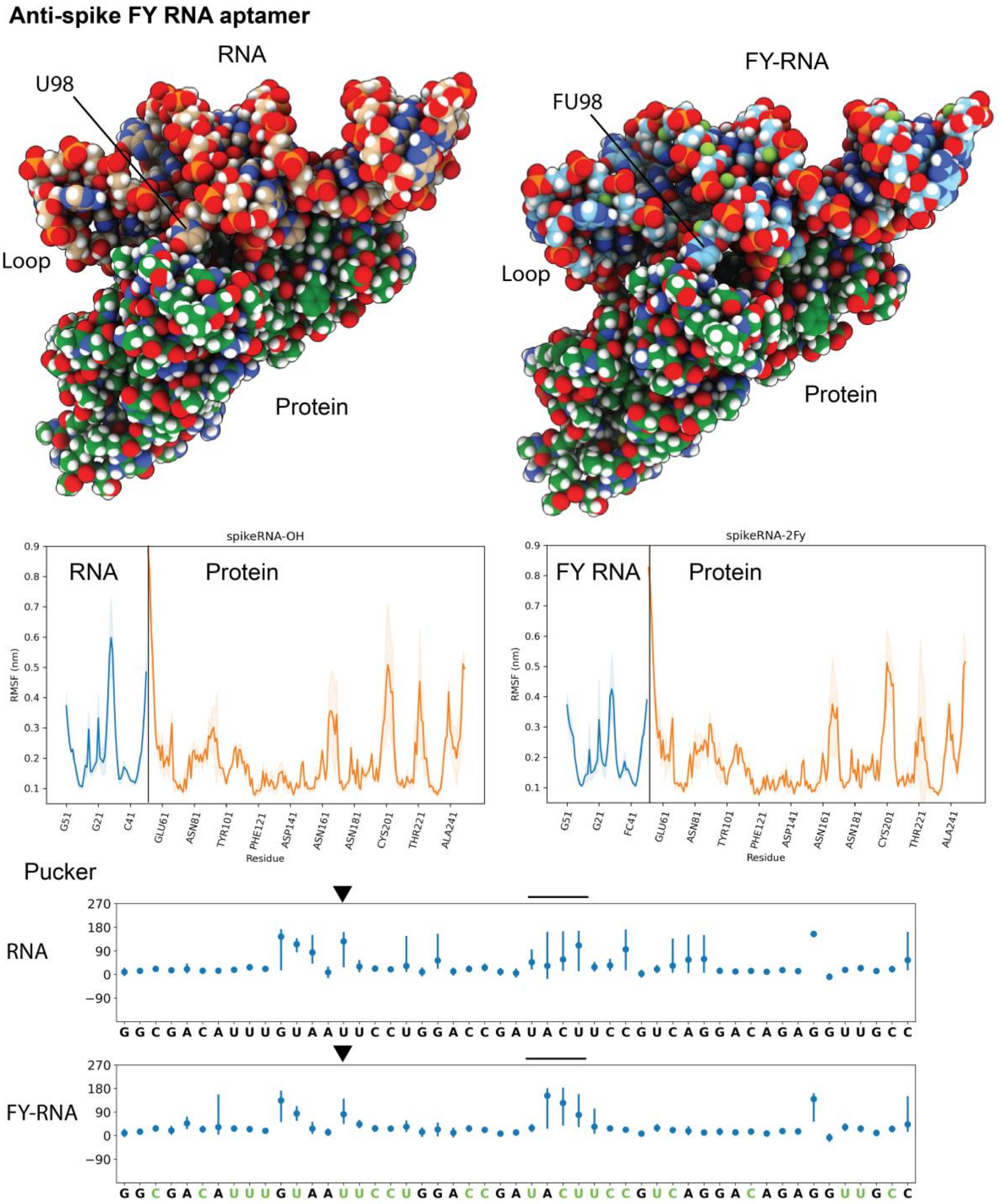

**Figure S21. MD-simulation of anti-Spike aptamer bound to the RBD.** Simulated structures of the anti-spike aptamer as RNA and FY-RNA bound to the RBD protein. Corresponding RMSF and Pucker plots are shown. The simulation reveals that fluctuation in the anti-spike aptamer was highest at the loop region and at specific residues (discussed in the main text). This fluctuation was particular high for RNA. Pucker plots with arrow heads showing the important residue U98 in the aptamer. Line indicated the loop region.

**Supplementary Table 1. RNA sequences.**

>PXT
GGAAUAUCGUCAUGGUGAUUCGUCACCAUGAGGCUAGAUCUCAUAUCUAGCGCUUUCGAGCGCUAGAGUCCUUAU
CUAGCCGGUUUAUACUUUCGAGUGUGAACCCGAUAUUCCGCGGAUCACUAUGAGUCGUUCGCGGCUCAUAGUCCG
GCUCAAAGGACAUCAUGGCCUGUUCGCAGGUUGUGAUUAUGAGUGAGCCGGGUAAGGCAUACCGUUCGCGGUAUG
UCUUACGAUCCGC

>5HT-TC
GGGCACUUACCCUUUAGUGCGAAGGUUUCGACCUUCGAUCCAUUUGUUCGCAAAUGGGAUGAUCUUCGGAUCAUC
CGCGUAGUCUGUUCAGUCGUUUCGACGACUGGCCCCACUUCGGUGGGGCCACGGUACUUAGAAGUGAACACUAAG
UGUCGUGAACACCAUUUGGUUAACUGCUCAAACUAAAUGGUGAUGAGGGAAGGAAUGACCCUCAUCGGACUACGC
GAUCCGAGUGAUGGGAAUGGCUGACCCAUCGCUCGGCACUGGAGGGUGAGUGCCCCUCAUUCGCAUAAGGGCCGA
CCCAGACAACAGCCAAGUUUGGGUCGGAGAUGCGAACAUUCCACGCAUCUGAACGGUUGAGAACUUACAAGGGCA
AGAGCAGAGUCCUUGUAAGGGCUUUACACGUCAAGUUCACAGACGUGUAAGGCCCGUCGCCCUUCGGGGCGACGU
UCACGGCAUUUCGAUGCCGUGCAGCCUGUUCGCAGGCUGCUUGACCGUUCCCCUGCCCUUUCGAGGGCAGACUAC
UCUUCGGAGUAGUCUUAUGUGAAUGAG

>6HBC
GGGAGAGUACUAUUCAGAUGCAGACCGCAAGUUCAGAGCGGUUUGCAUCUAGGGUACGUUUUCGAACGUAUCCUC
CGACUAAGUGUAUUCGUAUACUUAGUGCCUUGUGCCUGCUUCGGCAGGCAUGACCCAAAUGUGCCUUUCGGGGCA
CAUUUCCGGUCAUCCAAGUUCGCUUGGGUGAUGCGGGCGUAUAGGUUCGUCUAUACGUCCGCGUUUUCCGAGAAG
AGGUAACUCGGGAAACCGGUCCACGUGACAAAGGUAGAGUUACGUGGAGGGAGCAGCUGCAAAGGGAUAAUGCAG
UUGCUGGCUGGAUGCCAGAACUCACGACUGGCAUCUACGGGGAUGGUGCUCUCCCAAUUCUCCAUUUACCGCCGA
AUCGACCCCAACGUGAGAGGGGUCGGUUCCCCGAGCAUAGACCAAUAUCCCAGGUUUAUGCUCCCCAACGCUGGA
CGAACUACCUACGUCUAGCGUUCCGGCAAAUGAGUCAAUACCUCAGACUUAUUUGCGGUGCCUGAGCCUAAACUG
AACAUGGGUUCAGGCAUCUUGGCUCCAGUUCGCUGGAGCCGACGGUAGCGCUGCGUUCGCGCAGUGCUAGGGAGC
AUCCGUUUUCGAGCGGAUGCUGGGCGGUUGCCUGUUCGCAGGCAAUCGGGCCUACUCAUGAUUCGUCAUGAGUGG
UGACAGCGUGAUGUUCGCAUUACGCUGUCGGGUAGAUGGAGAAUU

>3HT-BP
GGAUACGUCUACGCUCAGUGACGGACUCUCUUCGGAGAGUCUGACAUCCGAACCAUACACGGAUGUGCCUCGCCG
AACAGUCUACGGCGAGCUUAAGCGCUGGGGACGCCCAACGCAUCACAAAGACUGAGUGAUGAACCAGAAGUAUGG
ACUGGUUGCGUUGGUGGAGACGGUCGGGUCCAGUUCGCUGUCGAGUAGAGUGUGGGCUCCAUCGACGCCGCUUUA
AGGUCCCCAAUCGUGGCGUGUCGGCCUGCUUCGGCAGGCACUGGCGCCGGGACCUUGAAGAGAUGAGAUUUCGAU
CUCAUCUUUGGGUGUCUCUGGUGCUUGAGGGCCCUGUGUUCGCACAGGGCCGCUCACUGGGUGUGGACGUAUCC

>Traptamer
GGAAUCUCGCCCGAUGUUCGCAUCGGGAUUUGCAGGUCCAUGGAUUACACCAUGCAACGCAGACCUGUAGAUGCC
ACGCUAGCCGUGGUGAGGGUCGGGUCCAGAUGUCAUUCGACUUUAACGCGCCUAAGCGUUGAAGGCGUGUUAGAG
CAGAUAGUUCGCUAUCUGGGGAGCCUGUUCGCAGGCUCAGGAGCCUUCGGGCUCCUAGCGCUAUUACCCCGGACA
CCACCGGGCAGACAAGUAAUGGUGCUCCUCGAAUGACUUCUGUUGAGUAGAGUGUGGGCUCCGCGGCUAGUGUGC
ACCUUAGCGGUGAAUGUCUGACACCGUUAAGGUGGUUACUCUUCGGAGUAACGCCGAGAUUCC

>PXT-antispike
GGAAGGUGGGGCACUUCGUUCGCGAAGUGCCGGCAAGAUCCCUCAUCUUGUACGUUCGCGUGCAAGACCAUCCAU
CUUGCCCAGGCGACAUUUGUAAUUCCUGGACCGAUACUUCCGUCAGGACAGAGGUUGCCUGCCACCUUCCACGAC
ACGGAAAGGGCUAUUCGUAGUCCUUUCCCUUCGUCAGGAUGGAUCUACCGGUGUUCGCAUCGGUAGAUUGAGGGU
GACGAAGAAUAUCGGAGGUGUUCGCGCCUUCGAUGUUGUGUCGU

**Supplementary Table 2. Cryo-EM data collection, refinement and validation statistics for FY-RNA PXT and 5HT.**

| | #1 FY-RNA PXT (J334) (EMD-53787) (PDB 9R7Q) | #2 FY-RNA 5HT (J225) (EMD-53795) (PDB 9R7W) |
|---|---|---|
| **Data collection and processing** | | |
| Magnification | 130,000x | 130,000x |
| Voltage (kV) | 300 | 300 |
| Electron exposure (e–/Å$^2$) | 60 | 60 |
| Defocus range (μm) | -2.0 to -0.8 | -2.0 to -0.8 |
| Pixel size (Å) | 1.2 | 1.2 |
| # mics. | 4,449 | 4,121 |
| Symmetry imposed | Non | Non |
| Initial particle images (no.) | 2,200,102 | 1,516,345 |
| Final particle images (no.) | 434,756 | 314,600 |
| Map resolution (Å) | 4.41 | 6.06 |
| FSC threshold | (0.143) | (0.143) |
| Map resolution range (Å) | 3.934-9.431 | 5.344-13.823 |
| **Refinement** | | |
| Initial model used (PDB code) | 8BTZ | 7PTS |
| Model resolution (Å) | 4.4 | N.A. |
| FSC threshold | (0.143) | |
| Model resolution range (Å) | 4.0/4.4/6.1 (0/0.143/0.5) | N.A. |
| Map sharpening *B* factor (Å$^2$) | 150.2 | 329.1 |
| Model composition | | |
| Non-hydrogen atoms | 5050 | 11752 |
| Nucleotide residues | 238 | 552 |
| Ligands | 0 | 0 |
| *B* factors (Å$^2$) | | |
| Nucleic acid | 80.06/874.00/214.30 (min/max/mean) | 119.81/873.54/343.46 (min/max/mean) |
| R.m.s. deviations | | |
| Bond lengths (Å) | 0.003 | 0.002 |
| Bond angles (°) | 1.111 | 1.048 |
| Validation | | |
| MolProbity score | 3.07 | 2.91 |
| Clashscore | 26.89 | 18.43 |

**Supplementary Table 3. Cryo-EM data collection, refinement and validation statistics for FY-RNA 6HBC.**

| | #3 FY-RNA 6HB-C dimer (J174) (EMD-54779) (PDB 9SD9) | #4 FY-RNA 6HB-C dimer (focus) (J280) | #5 FY-RNA 6HB-C mono. (J49) (EMD-53803) (PDB 9R82) |
|---|---|---|---|
| **Data collection and processing** | | | |
| Magnification | 130,000x | 130,000x | 130,000x |
| Voltage (kV) | 300 | 300 | 300 |
| Electron exposure (e–/Å$^2$) | 60 | 60 | 60 |
| Defocus range (μm) | -2.0 to -0.8 | -2.0 to -0.8 | -2.0 to -0.8 |
| Pixel size (Å) | 1.2 | 1.2 | 1.2 |
| # mics. | 3,538 | 3,538 | 3,538 |
| Symmetry imposed | Non | C2 | Non |
| Initial particle images (no.) | 87,516 (Topaz) | 87,516 (Topaz) | 746,064 |
| Final particle images (no.) | 49,938 | 62,995 (sym. exp.) | 46,693 |
| Map resolution (Å) | 9.6 | 9.0 | 8.12 |
| FSC threshold | (0.143) | (0.143) | (0.143) |
| Map resolution range (Å) | 8.546-15.913 | 8.049-13.588 | 7.749-17.945 |
| **Refinement** | | | |
| Initial model used (PDB code) | N.A. | N.A. | N.A. |
| Model resolution (Å) | N.A. | | N.A. |
| FSC threshold | | | (0.143) |
| Model resolution range (Å) | N.A. | | N.A. |
| Map sharpening *B* factor (Å$^2$) | N.A. | N.A. | N.A. |
| Model composition | | | |
| Non-hydrogen atoms | 30671 | | 15523 |
| Nucleotide residues | 1440 | | 728 |
| Ligands | 0 | | 0 |
| *B* factors (Å$^2$) | | | |
| Nucleic acid | 166.38/1064.10/544.62 (min/max/mean) | | |
| R.m.s. deviations | | | |
| Bond lengths (Å) | 0.001 | | 0.003 |
| Bond angles (°) | 0.902 | | 1.191 |
| Validation | | | |
| MolProbity score | 2.76 | | 3.04 |
| Clashscore | 12.43 | | 24.90 |

**Supplementary Table 4. Cryo-EM data collection statistics for FY-RNA 3HT-BP and Traptamer.**

| | #6 FY-RNA 3HT-BP (J214) (EMD-55626) | #7 FY-RNA Traptamer (J45) (EMD-55635) |
|---|---|---|
| **Data collection and processing** | | |
| Magnification | 130,000x | 130,000x |
| Voltage (kV) | 300 | 300 |
| Electron exposure (e–/Å$^2$) | 60 | 60 |
| Defocus range (μm) | -2.0 to -0.8 | -2.0 to -0.8 |
| Pixel size (Å) | 1.2 | 1.2 |
| # mics. | 8,642 | 4,121 |
| Symmetry imposed | Non | Non |
| Initial particle images (no.) | 1,754,761 | 19,111 (Topaz) |
| Final particle images (no.) | 107,281 | 7,214 |
| Map resolution (Å) | 6.42 | 9.93 |
| FSC threshold | (0.143) | (0.143) |
| Map resolution range (Å) | 5.550-13.628 | 8.920-17.344 |

**Supplementary Table 5. Cryo-EM data collection, refinement and validation statistics for FY-RNA PXT-Spike aptamer and Spike protein.**

| | #8 FY-RNA PXT Spike trimer (J586) (EMD-55623) | #9 Spike RBD +aptamer (focus) (J620) (EMD-55625) (PDB 9T74) | #10 Spike NTD (focus) (J598) (EMD-55624) | #11 Spike core (focus) (J670) (EMD-55622) | #12 FY-RNA PXT (focus) (J687) (EMD-55621) |
|---|---|---|---|---|---|
| **Data collection and processing** | | | | | |
| Magnification | 130.000x | 130.000x | 130.000x | 130.000x | 130.000x |
| Voltage (kV) | 300 | 300 | 300 | 300 | 300 |
| Electron exposure (e–/Å$^2$) | 60 | 60 | 60 | 60 | 60 |
| Defocus range (μm) | -2.0 to -0.8 | -2.0 to -0.8 | -2.0 to -0.8 | -2.0 to -0.8 | -2.0 to -0.8 |
| Pixel size (Å) | 0.971 | 0.971 | 0.971 | 0.971 | 0.647 |
| Symmetry imposed | no | C3 | C3 | C3 | C1 |
| Initial particle images (no.) | 875,298 | 875,298 | 875,298 | 875,298 | 1,506,092 |
| Final particle images (no.) | 433,417 | 736,847 (sym. exp.) | 794,657 (sym. exp.) | 1,300,251 (sym. exp.) | 109,814 |
| Map resolution (Å) FSC threshold | 2.90 (0.143) | 3.46 (0.143) | 3.02 (0.143) | 2.78 (0.143) | 6.92 (0.143) |
| Map resolution range (Å) | 2.490 – 45.259 | 2.083 - 48.822 | 2.387 - 46.248 | 2.342 - 46.369 | 6.581 - 13.878 |
| **Refinement** | | | | | |
| Initial model used (PDB code) | | N.A. | | | |
| Model resolution (Å) FSC threshold | | 3.4 (0.143) | | | |
| Model resolution range (Å) | | 3.3/3.4/3.5 (0/0.143/0.5) | | | |
| Map sharpening *B* factor (Å$^2$) | | 154.3 | | | |
| Model composition | | | | | |
| Non-hydrogen atoms | | 2673 | | | |
| Nucleotide residues | | 51 | | | |
| Protein | | 197 | | | |
| Magnesium | | 1 | | | |
| NAG | | 2 | | | |
| *B* factors (Å$^2$) | | | | | |
| Nucleic acid | | 50.27/145.63/94.28 | | | |
| Protein | | 10.46/92.43/38.98 | | | |
| Ligant (Mg) | | 0.00/79.33/61.07 (min/max/mean) | | | |
| R.m.s. deviations | | | | | |
| Bond lengths (Å) | | 0.025 (51) | | | |
| Bond angles (°) | | 2.532 (51) | | | |
| Validation | | | | | |
| MolProbity score | | 2.18 | | | |
| Clashscore | | 5.53 | | | |

**Supplementary Table 6. Lifetime fluorescence measurements.**

| | **RNA** | **FY RNA** |
|---|---|---|
| **Lifetime of DFHBI-1T Broccoli aptamer** | 4.583 ± 0.005 ns | 4.564 ± 0.001 ns |
| **Lifetime of HBC620 Pepper aptamer** | 3.97 ± 0.04 ns | 4.117 ± 0.005 ns |

The data was obtained by fitting fluorescence lifetime measurements by a monoexponentional decay function. The data is noted as the mean ± standard deviation (n=3).

**Supplementary Video 1. MD simulation of PXT as RNA and FY-RNA.** The video shows that the helices of TL1 and TL3 (bottom) tend to interact and become parallel in the RNA PXT (left), whereas this is not observed for the FY-RNA PXT (right).

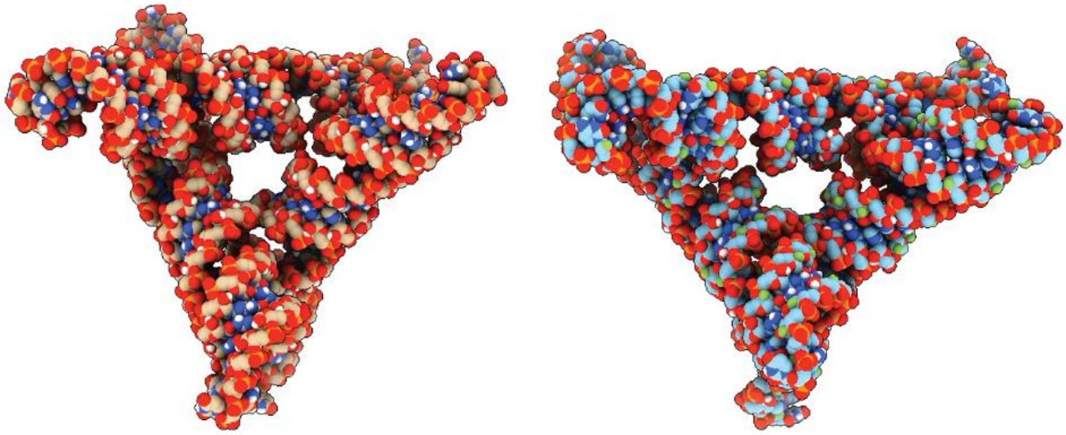

**Supplementary Video 2. MD simulation of 5HT as RNA and FY-RNA.** The video shows that the 5HT is more compact as RNA (left) and less compact as FY-RNA (right), which is likely due to less H-bond interactions between the helices.

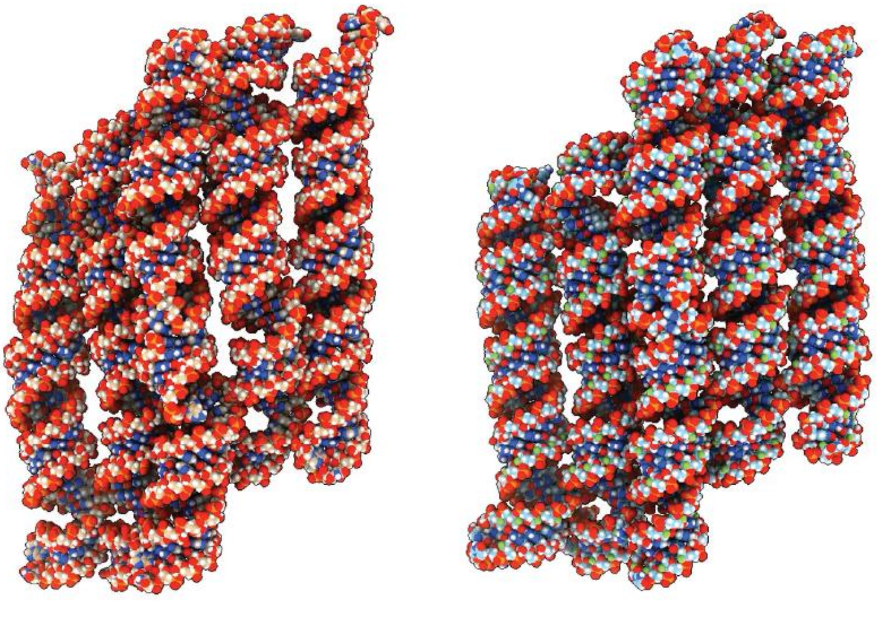

**Supplementary Video 3. MD simulation of the X-90 crossover.** Simulated structures of the X-90 as RNA, FY-RNA and F-RNA (from left to right). The movie shows that RNA helices align in parallel, FY-RNA helices are more dynamic, and F-RNA helices make a stable conformation.

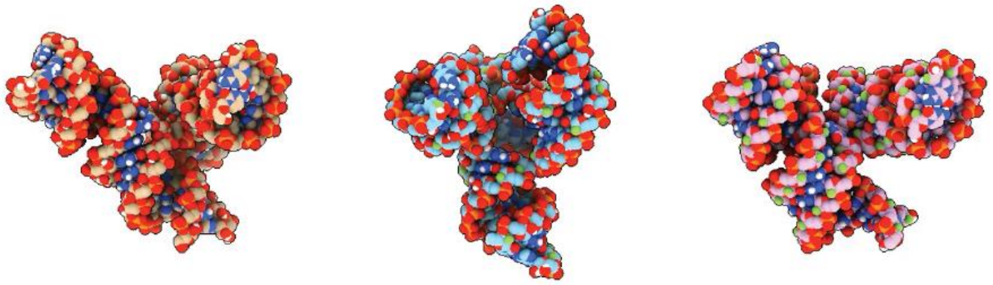

**Supplementary Video 4. MD simulation of anti-Spike aptamer bound to the RBD.** Simulated structures of the anti-spike aptamer as RNA (left) and FY-RNA (right) bound to the RBD protein. The movie shows that the tetraloop is more flexible and the U98 does to keep a stable stacking with Tyr505 as RNA.

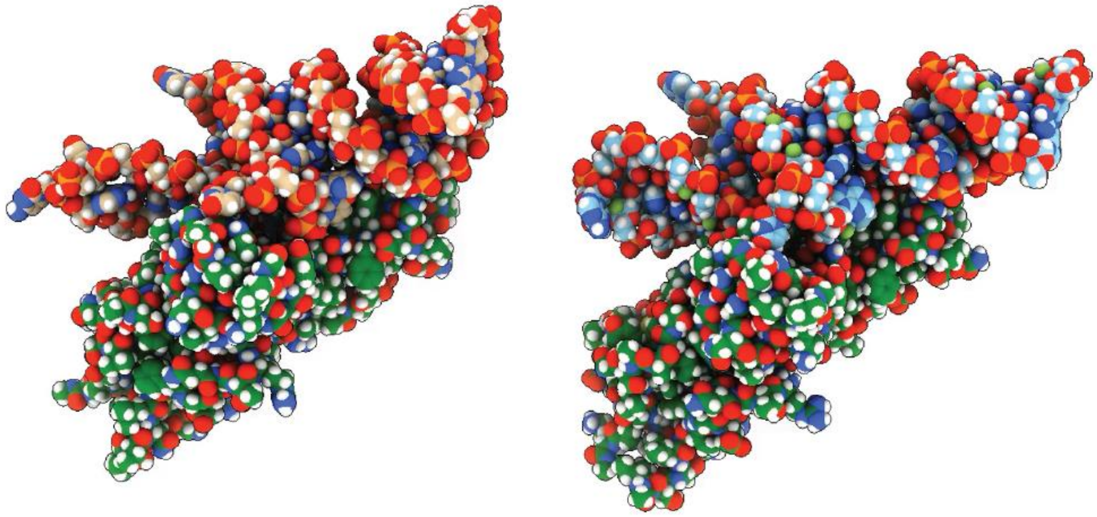

**Supplementary Video 5. Detail of the MD simulation showing the FU98-Tyr505 interaction.** The movie shows that the FU98-Tyr505 interaction is maintained during the MD simulation of the FY-RNA anti-Spike aptamer bound to the RBD.

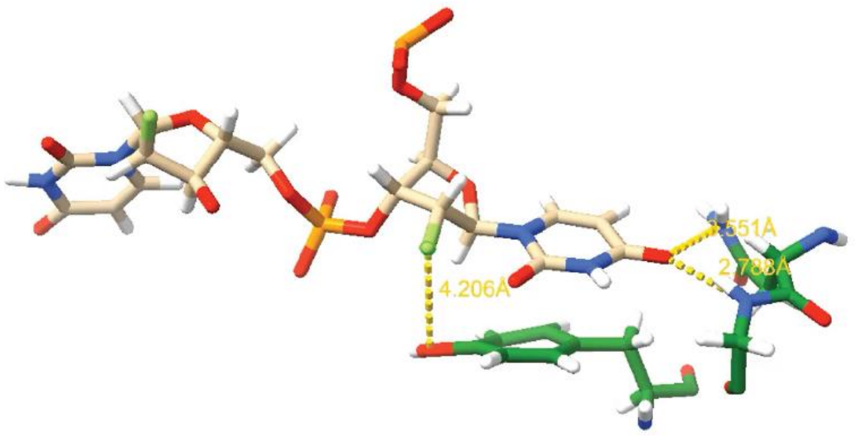

**Supplementary Video 6. Detail of the MD simulation showing the U98-Tyr505 interaction.** Movie shows that the U98-Tyr505 interaction is maintained during the MD simulation of the anti-Spike aptamer bound to the RBD, when changed to RNA.

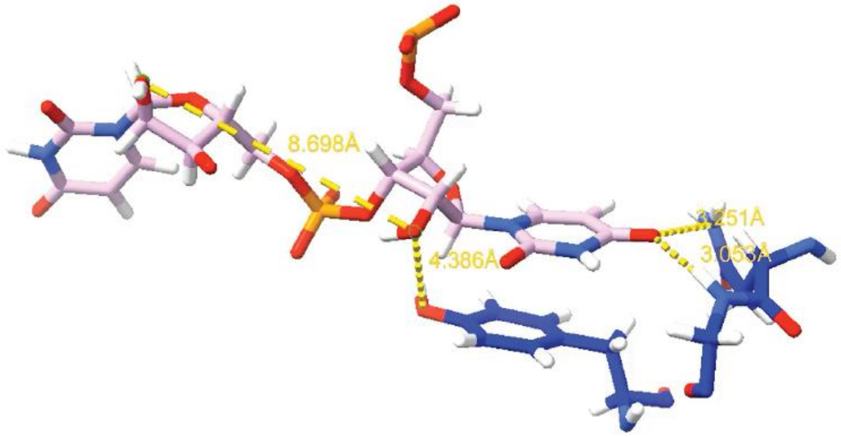
